# Computational modeling of neurotransmitter cycling predicts human brain glutamate and GABA dynamics in response to administration of exogenous ketones

**DOI:** 10.64898/2026.02.11.700015

**Authors:** Botond B. Antal, Lilianne R. Mujica-Parodi, Helmut H. Strey, Eva-Maria Ratai, Silvia Mangia, Douglas L. Rothman

**Affiliations:** Center for Magnetic Resonance Imaging, University of Minnesota, Minneapolis, MN, USA; Department of Biomedical Engineering, Stony Brook University, Stony Brook, NY, USA; Computer Science and Artificial Intelligence Laboratory, Massachusetts Institute of Technology, Cambridge, MA, USA; Athinoula A. Martinos Center for Biomedical Imaging, Department of Radiology, Massachusetts General Hospital, Harvard Medical School, Charlestown, MA, USA; Laufer Center for Physical and Quantitative Biology, Stony Brook University, Stony Brook, NY, USA; Santa Fe Institute, Santa Fe, NM, USA; Magnetic Resonance Research Center and Departments of Radiology and Biomedical Engineering, Yale University, New Haven, Connecticut, USA

## Abstract

Administration of ketones is used as a therapeutic option in multiple conditions, including epilepsy, mental health disorders, and brain aging. A proposed mechanism of action involves the modulation of glutamate and GABA, the brain’s primary excitatory and inhibitory neurotransmitters, which jointly regulate the excitatory-inhibitory balance. However, the precise mechanism by which ketones influence these neurotransmitters remains unclear. In this study, we hypothesize that ketones modulate glutamate and GABA through the pseudomalate-spartate shuttle (PMAS). To test this, we developed a computational model of neurotransmitter cycling centered on the PMAS, simulating the temporal dynamics and steady-state concentrations of glutamate and GABA as functions of ketone metabolism. We then compared the model outputs with MRS data from ketone administration experiments and found agreement with the model predictions, providing quantitative support for the model. Building on this agreement, we performed metabolic control analysis, which identified partial displacement of glucose metabolism through the PMAS as the dominant mechanism underlying the observed reductions in glutamate and GABA and revealed key enzymes that selectively modulate each neurotransmitter. Overall, the model provides researchers and clinicians with a framework for hypothesis testing and treatment optimization, while also serving as a foundation for future model expansions.

## 1 Introduction

Ketone bodies are alternative energy substrates to glucose with therapeutic relevance in brain disorders. Ketogenic therapies are already clinically established for refractory epilepsy [1], and recent studies suggest potential benefits in mental health disorders [2], age-related neurodegeneration [3], and even in improving cognitive and functional biomarkers in healthy aging [4, 5]. Positron emission tomography (PET) [6] and ^13^C magnetic resonance spectroscopy (^13^C MRS) [7] studies have demonstrated that ketones are taken up and utilized by the human brain as fuel, whether produced endogenously or administered exogenously. Beyond their role as an energy substrate, ketones may also exert neuromodulatory effects by altering cellular levels of glutamate and gamma-aminobutyric acid (GABA), the primary excitatory and inhibitory neurotransmitters in the brain, with changes in intracellular concentrations directly propagating to the vesicular and extracellular pools via glutamate and GABA membrane transporters [8, 9].

Consistent with a role of glutamate and GABA in the neuromodulatory effects of ketones, multiple studies have reported changes in brain levels of glutamate and GABA in chronic ketosis [10, 11], and more recently, an MRS study with a mechanistic focus on short time-scales has shown that acute administration of *β*-hydroxybutyrate reduces glutamate and GABA levels by 11% and 37%, respectively [12, 13]. These results have important implications for understanding the mechanisms underlying the clinical benefits of ketone-based therapies and for their optimization, because the dysregulation of glutamate and GABA levels is a hallmark of various brain disorders, including age-related neurodegenerative diseases [14–16] and psychiatric conditions [17–19].

Despite their therapeutic potential, the lack of consensus regarding the mechanisms by which ketones modulate neurotransmitters remains a challenge for their therapeutic application. It is established that the maintenance of glutamate and GABA pools, which are approximately 80% recycled and 20% synthesized de novo [20, 21], shares overlapping energy metabolic pathways with ketone oxidation [4]. Furthermore, neurotransmitters facilitate synaptic activity, a set of processes which carry a large energy cost, estimated to be 41% of total ATP use in the cortex [22, 23].

Identifying the mechanisms by which ketone administration modulates brain metabolism and the excitatory/inhibitory balance requires integrating diverse experimental observations spanning ketone pharmacokinetics, neurotransmitter metabolism, and established energetic constraints. This integration is particularly challenging because of the presence of three major cell types with highly specialized metabolic functions: glutamatergic neurons, GABAergic neurons, and glial cells. These compartments are coupled through neuronal-glial trafficking, in which astrocytes regulate interstitial neurotransmitter concentrations by taking up neuronally released glutamate and GABA, converting them to glutamine, and subsequently returning them to neurons through the glutamate-glutamine and GABA-glutamine cycles. To address this complexity, several computational models incorporating separate metabolic compartments for glutamatergic neurons, GABAergic neurons, and glia have been developed [24–27]. However, a limitation of existing approaches is that pathway fluxes, metabolite concentrations, and enzyme kinetic properties are largely derived from in vitro animal studies and estimates of enzyme activity based on gene expression data. Because in vitro measurements can be affected by postmortem autolysis and may differ substantially from in vivo conditions, these parameter values may not accurately reflect metabolism in the living human brain [28, 29].

We describe here the development and application of a computational metabolic model and its use to determine the mechanistic basis of the changes in glutamate and GABA metabolism during ketone exposure. The model combines the pharmacokinetics of ketone utilization, from transport across the blood-brain barrier to subsequent oxidation, with neurotransmitter cycling represented through enzymatic reactions in neurons and astrocytes. To describe these processes, we integrated algebraic expressions and differential equations into a unified framework, with parameters derived primarily from in vivo studies. The model was then constrained by the experimental findings of a close to stoichiometric 1:1 coupling between neurotransmitter cycling rates and oxidative glucose metabolism [20, 23, 30, 31]. Recent metabolic modeling work showed that this stoichiometry derives from a pathway referred to as the pseudo-malate-aspartate shuttle (PMAS) [32], which connects the glycolytic steps of glucose oxidation to the mitochondrial conversion of glutamine to glutamate, a key reaction in neurotransmitter cycling. The PMAS may therefore represent a key site where ketones exert their effects on glutamate and GABA, as PET studies have shown that ketones can partially displace glucose metabolism even when glucose is sufficiently available [33, 34].

We then tested whether the framework could reproduce the experimentally observed reductions in glutamate and GABA [13] when driven by the corresponding experimental blood ketone measurements. We found that, without modification, the model predicted the measured decreases in glutamate and GABA. Using metabolic control analysis as a form of sensitivity analysis, we identified the primary mechanism as *β*-hydroxybutyrate partially displacing glucose metabolism and thereby reducing flux through the PMAS [32], slowing neurotransmitter cycling and reducing the steady-state concentrations of glutamate and GABA. The greater percentage decrease in GABA was attributed to the combination of PMAS inhibition by ketone metabolism and the kinetic properties of the enzymes involved in GABA synthesis (glutamate decarboxylase 65 and 67) and breakdown (GABA transaminase). Although applied specifically to acute ketone infusion, the model’s metabolic network equations, stoichiometric relationships, and enzyme and transporter kinetics were determined independently. Therefore, the model holds promise for interpreting and guiding pharmacological and metabolic interventions in the brain.

## 2 Methods

### 2.1 Overview of modeling neurotransmitter cycling

The dynamics of glutamate and GABA were modeled based on the reactions of neurotransmitter cycling within a connected system of glutamatergic neurons, GABAergic neurons, and astrocytes (Figure 1A). This framework was motivated by prior studies showing that glutamate and GABA are released by neurons, taken up by astrocytes, converted to glutamine, and subsequently cycled back to neurons [35]. Specifically, ^13^C MRS experiments indicate that approximately 80% of glutamate is directly converted to glutamine, while the remainder is first oxidized and then resynthesized to glutamate and glutamine [36]. Additionally, the model was further informed by the established enzymatic processes unique to GABAergic neurons that govern the synthesis, release, and cycling of distinct GABA pools (Figure 1B).

**Fig. 1.**
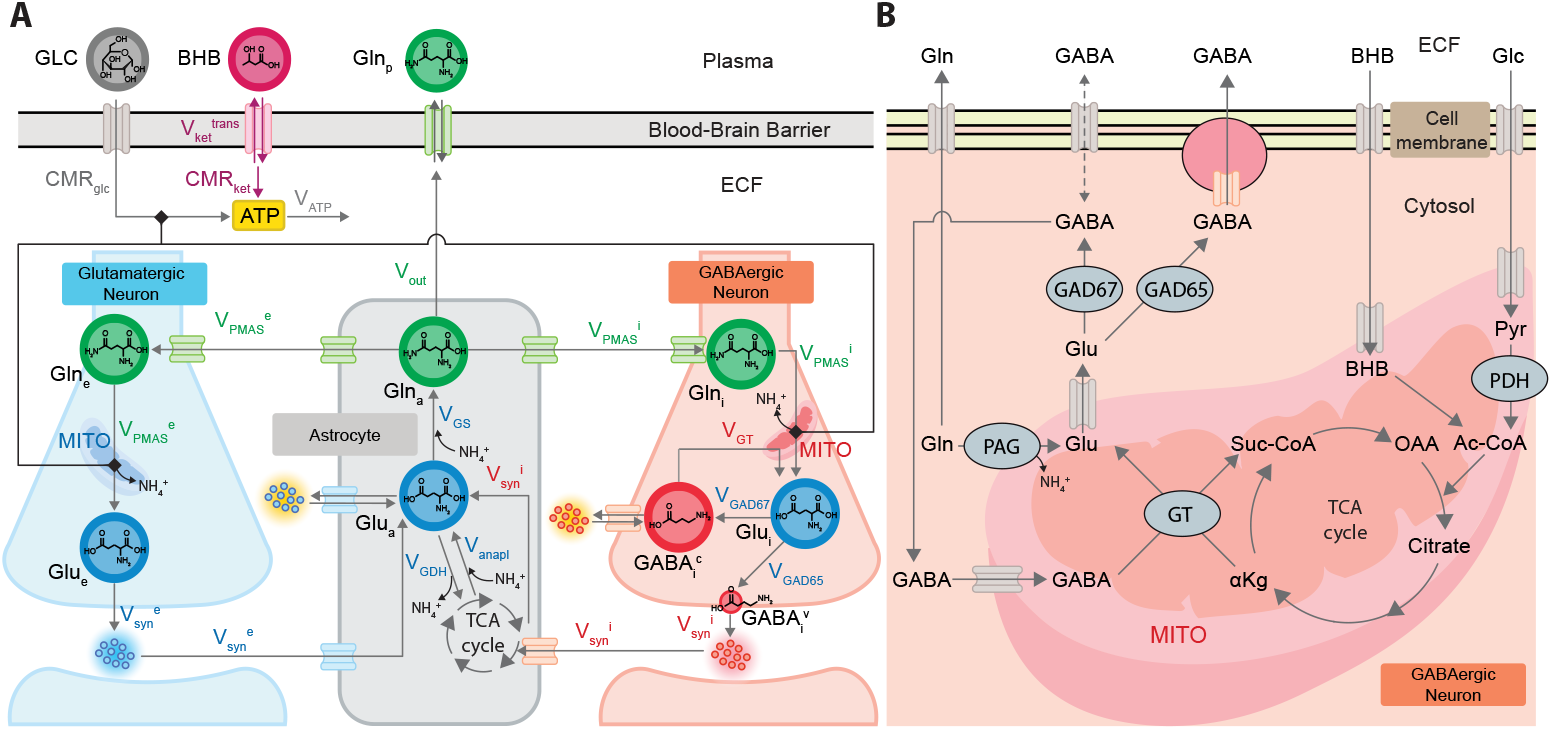
Overview of the computational model and related biochemical pathways. **A**: Schematic representation of the model. The two energy substrates, glucose (GLC) and D-*β*HB (BHB), are shown at the top, with arrows indicating transport and metabolic utilization. D-*β*HB partially displaces glucose metabolism, which is coupled to neurotransmitter cycling via the pseudomalate–aspartate shuttle (PMAS; black lines with square arrowheads). Blue, green, and red disks with black labels depict the major glutamate (Glu), glutamine (Gln), and GABA pools represented in the model (e: excitatory, i: inhibitory, a: astrocytic, c: cytosolic, v: vesicular, p: plasma). Arrows with colored labels denote reactions for which kinetics were explicitly modeled. Particles with yellow backgrounds indicate neuromodulatory pools of glutamate and GABA that are in equilibrium with the intracellular glutamate pool in astrocytes and the cytosolic GABA pool in GABAergic neurons, respectively. These pools are distinct from their synaptic counterparts (particles marked with blue and red backgrounds) and exert neuromodulatory effects that are different from classical neurotransmitter-dependent synaptic signaling. **B**: Detailed biochemical pathways related to the modeled processes in the GABAergic neuron. Of particular significance are the GABA shunt, catalyzed by GABA-T, the bifurcation of GABA synthesis fluxes catalyzed by GAD65 and GAD67, and the competition between GLC and BHB as energy substrates to the oxidative apparatus. Abbreviations: GS, glutamine synthetase; GT, GABA transaminase; GAD65, glutamate decarboxylase 65; GAD67, glutamate decarboxylase 67; GDH, glutamate dehydrogenase; anapl, anaplerosis; syn, synaptic signaling; mito, mitochondrion; ECF, extracellular fluid; PDH, pyruvate dehydrogenase; Ac-CoA: acetyl coenzyme A; *α*Kg, alpha-ketoglutarate; pyr, pyruvate; OAA, oxaloacetate; Suc-CoA, succinyl coenzyme A.

While neurotransmitter cycling involves many reactions, we applied established principles from metabolic control analysis (MCA) to reduce the number of reactions that require explicit representation for accurate simulations. According to MCA, although control of pathway fluxes and metabolite concentrations is distributed throughout the metabolic network, a subset of enzymes and transporters typically exerts most of the control based on their location in the network and their kinetic properties [37]. For example, the first step in a pathway usually exerts the majority of flux control, whereas reversible reactions in the middle of pathways often have low control and can therefore be modeled as flow-throughs. In the following sections, we describe in detail the kinetics and corresponding parameters of the reactions that were explicitly included in the model.

To simulate the system dynamics, we used differential equations for reactions whose kinetics evolve on the timescale of interest, and algebraic expressions for reactions that catalyze rapid exchanges between substrates and products, effectively keeping them near equilibrium values, as well as for reactions that, due to high enzyme activity, do not influence flux. In addition, we took advantage of stoichiometric relationships between fluxes across membranes separating specific compartments (e.g., mitochondrial–cytosolic and neuronal–glial) that are enforced by the stoichiometry of the transporters involved [32]. These relationships strongly constrain the relative fluxes, allowing a more compact representation of the system without loss of mechanistic detail.

Summaries of parameter values and concentrations are provided in Table 1 and Table 2. The references used to estimate the compartmental concentrations are summarized in SI Table 1, while the sources for parameter values are described in the main text. All concentrations originally reported in *µ*mol/g were converted to mM assuming a brain tissue density of 1 g/ml. Concentrations and reactions are expressed separately for each compartment and are denoted by subscripts *e* for excitatory (i.e., glutamatergic) neurons, *i* for inhibitory (i.e., GABAergic) neurons (which together account for approximately 90% of cortical synapses [38]), and *a* for astrocytes. We applied a fixed volumetric composition throughout the study to convert between total tissue and compartmental concentrations. Tissue volume was partitioned into neurons (70%), astrocytes (20%), and extracellular space (10%) [39, 40]. The neuronal fraction (70%) was subdivided assuming a 4:1 glutamatergic:GABAergic ratio [41] corresponding to 56% and 14% of total tissue volume for glutamatergic and GABAergic neurons, respectively.

**Table 1.**
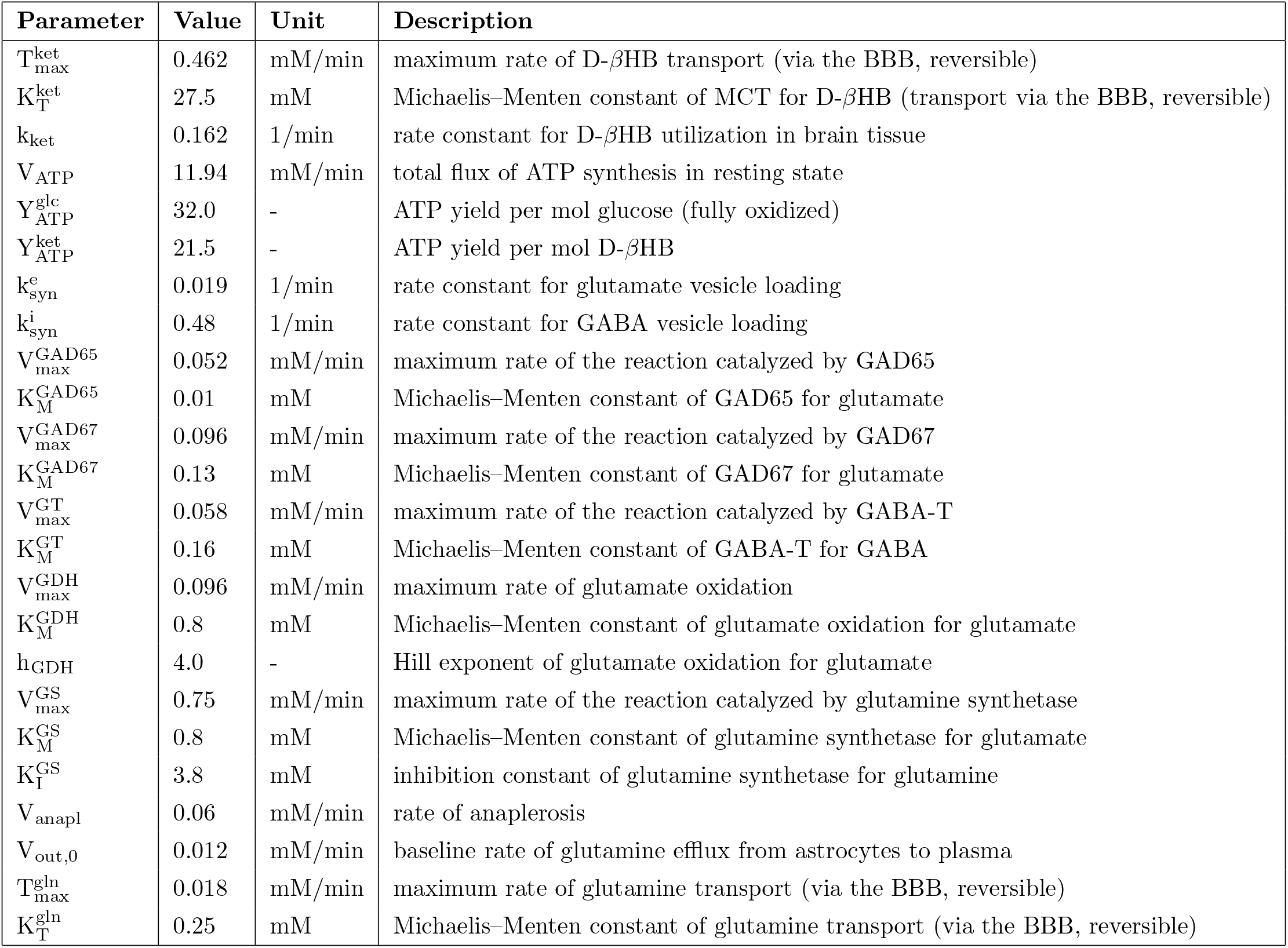
Parameter values used in the model equations.

**Table 2.** Steady-state metabolite concentrations used as initial conditions in simulations.

| Metabolite | Value (mM) | Description |
| --- | --- | --- |
| [Glu]<sub>e</sub> | 10.0 | Glutamate pool of glutamatergic neurons |
| [Glu]<sub>i</sub> | 0.13 | Glutamate pool of GABAergic neurons |
| [Glu]<sub>a</sub> | 0.8 | Glutamate pool of astrocytes |
| [Gln]<sub>e</sub> | 0.28 | Glutamine pool of glutamatergic neurons |
| [Gln]<sub>i</sub> | 0.07 | Glutamine pool of GABAergic neurons |
| [Gln]<sub>a</sub> | 3.8 | Glutamine pool of astrocytes |
| [GABA]<sub>i</sub><sup>c</sup> | 0.73 | Cytosolic GABA pool of GABAergic neurons |
| [GABA]<sub>i</sub><sup>v</sup> | 0.1 | Vesicular GABA pool of GABAergic neurons |
| [Ket]<sub>b</sub> | 0 | Brain tissue D-βHB |

The model assumes resting awake state, in which the subject (human or animal model) is not given focused external stimuli such as sensory stimulation or tasks to perform. The large majority of parameters in the literature have been determined under this condition, including human neuroimaging studies, which the specific parameters of our model were chosen to reflect. The experimental dataset, described later and used to test consistency with the model, was likewise acquired under a resting-state condition.

### 2.2 Coupling between neurotransmitter cycling and glucose metabolism

Prior work has shown that the glutamine-to-glutamate reaction catalyzed by phosphate-activated glutaminase (PAG) exerts strong flux control over neurotransmitter cycling [32]. Motivated by this, PAG was the first key reaction for which we assigned kinetics in our model. Because PAG operates within the PMAS [32], its rate was constrained to that of the overall PMAS. Although the PMAS encompasses several reactions, we modeled it as a single unit based on the argument that these reactions are tightly coupled by stoichiometry, as elaborated in [32].

We determined the value of *V*_*PMAS*_ based on two observations. First, the rate of PMAS governs neurotransmitter cycling at metabolic steady-state, such that *V*_*NTcyc*_ = *V*_*PMAS*_. Second, a prior report combining data from multiple species established the coupling between the neurotransmitter cycling flux and the cerebral metabolic rate of oxidative glucose metabolism in neurons 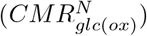 [29], as discussed in more detail later. On this basis, we express *V*_*PMAS*_ as a function of 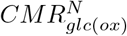.

For the calculation of 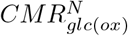, we first adopted a resting-state baseline value for total gray matter *CMR*_*glc*_ of 0.41 mM/min, based on aggregated results from previously published human PET and MRSI data [29]. As prior literature has demonstrated that resting-state *CMR*_*glc*_ values are relatively homogeneous (within *±*10% of means) across different brain regions [42], we considered the chosen value of *CMR*_*glc*_ to reasonably hold for any brain region. To derive 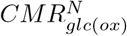 from *CMR*_*glc*_, first the oxidative portion was taken assuming an oxygen-glucose index (OGI) of 5.5 (91%) based on [43]:

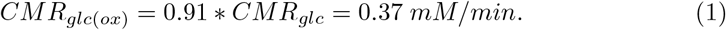

Next, we considered the neuronal fraction based on prior human studies indicating that the glial component is approximately 25% of the total *CMR*_*glc*_ [40], leaving 75% for the neuronal contribution. Accordingly:

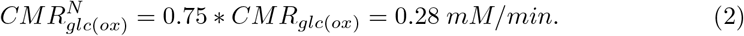

From 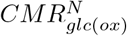, we then aimed to determine *V*_*PMAS*_ separately for both glutamatergic 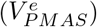 and GABAergic 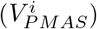 subpopulations, based on [29] showing that the in vivo coupling in animal models and human cerebral cortex between glucose oxidation and neurotransmitter cycling rates hold separately in both cell types. The same work has established that 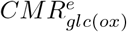 and 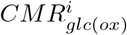 are approximately 83% and 17% of 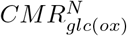, respectively. Based on this distribution, we expressed the population specific values with the following equations:

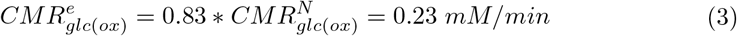

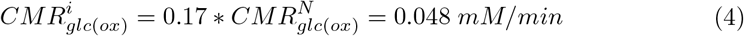

From 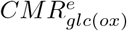 and 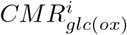 we then determined 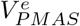 and 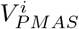 using the following formulas, also adapted from [29] and substituted in for steady-state:

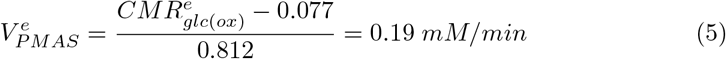

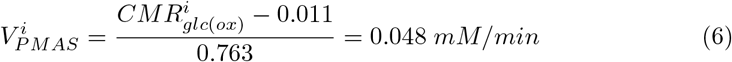

Based on 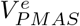, we described the flux of glutamate release through synaptic activity 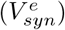. Prior work has shown that the rate of synaptic vesicle loading with glutamate scales approximately linearly with cytosolic glutamate availability ([*Glu*]_*e*_) [44], thus 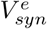 can be expressed using a rate constant 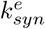:

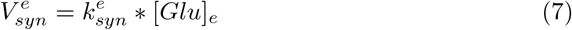

For steady-state, where 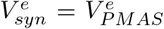, and a resting-state value for [*Glu*]_*e*_ = 10 mM [45–47], 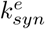 is then given by:

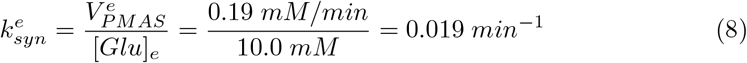

### 2.3 Kinetics of GABA cycling

GABA metabolism differs in key ways from glutamate metabolism and therefore warrants distinct kinetic descriptions. For glutamate, the general consensus is that there is a single dominant pool (*Glu*_*e*_), which interfaces with synaptic function. In contrast, evidence suggests the presence of two distinct GABA pools within GABAergic neurons, produced by two glutamic acid decarboxylase (GAD) isoforms from glutamate (*Glu*_*i*_) [48, 49] (Figure 1B). The first isoform, GAD65, is localized to vesicular membranes and axonal terminals and supplies the vesicular GABA pool responsible for synaptic signaling 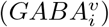, with its flux modulated by neuronal activity [50]. In contrast, the more homogeneously distributed isoform GAD67 produces cytosolic GABA 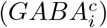 and exhibits activity that is insensitive to the GABA vesicular neurotransmitter release and cycling rate [50].

To model GABA kinetics, we first defined compartment-specific concentrations for the relevant metabolic pools. In vivo MRS studies reported a total GABA concentration of approximately 1 mM [51, 52], which we assumed to reside predominantly within GABAergic neurons. This value was corrected for homocarnosine, estimated to contribute approximately 17% to the measured GABA signal [53], yielding a total GABA concentration of 0.83 mM distributed across cytosolic and vesicular pools.

The vesicular GABA pool 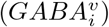 was estimated at 0.1 mM, with the remaining 0.73 mM assigned to the cytosolic pool 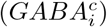. This partitioning was based on an intravesicular GABA concentration of 100 mM [54] and the assumption that GABAergic vesicles occupy approximately 0.1% of total tissue volume. The latter estimate was derived from reported synaptic densities of approximately 10^9^ synapses per mm^3^ [55], approximately 250 vesicles per bouton [56], a vesicle diameter of 40 nm [57], and the assumption that approximately 20% of synapses are GABAergic [58].

For setting [*Glu*]_*i*_, we considered ^13^C MRS studies showing no delay in GABA labeling relative to glutamate [59], which implies a relatively low intracellular glutamate concentration, as larger pools would be expected to introduce a measurable lag in GABA dynamics [60]. Accordingly, [*Glu*]_*i*_ was set to 0.13 mM, equal to the estimated Michaelis–Menten constant of GAD67 derived from in vitro measurements [61–63]. This value is also consistent with reports that GABAergic neurons contain approximately 2% of total glutamate [64], representing a substantially smaller fraction than in glutamatergic neurons.

Assuming that the majority of GABA involved in neurotransmitter cycling passes through synaptic signaling and 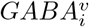, in homeostatic states, flux through GAD65 must maintain balance with the rate of 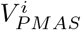. As such, in steady-state:

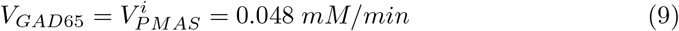

The GAD65 rate was modeled using Michaelis–Menten kinetics with low elasticity for glutamate (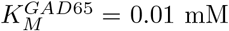 relative to [*Glu*]_*i*_ = 0.13 mM):

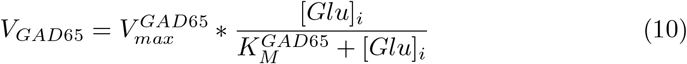

The low *K*_*M*_ of GAD65 was based on prior evidence showing that reductions in intracellular glutamate do not substantially affect synaptic GABA release, in contrast to the cytosolic GABA pool produced by GAD67, which is sensitive to such decreases [48, 50]. As a result, the synthesis rate of 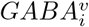 remains approximately constant across physiological [*Glu*]_*i*_ values in the resting state.

Given *V*_*GAD*65_ = 0.048 mM/min, 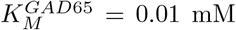, and [*Glu*]_*i*_ = 0.13 mM, 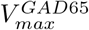 can be expressed and quantified from Equation 10, yielding a value of 0.052 mM/min. The resulting 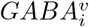 is then released during GABAergic synaptic activity at a rate analogous to 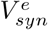, following Equation 7:

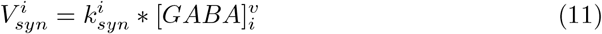

Where the value of 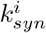 can be calculated in steady-state given 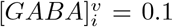 mM:

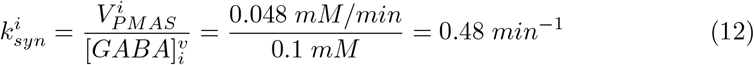

The cytosolic GABA pool 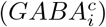, which constitutes the majority of total GABA in brain tissue, is primarily produced by GAD67. Rather than being released, 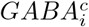 is cycled back to *Glu*_*i*_ via the GABA shunt, an intracellular cycling pathway specific to GABAergic neurons [65]. In the model, the GABA shunt was represented as two opposing reactions. The forward reaction corresponds to the conversion of *Glu*_*i*_ to *GABA*^*c*^ catalyzed by GAD67, with its rate described as:

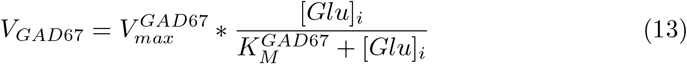

Based on estimates of the relative signaling and non-signaling fluxes in the resting awake state [29] and evidence suggesting that GAD67 accounts for approximately 50% of the total GABA synthesis and essentially all of the non-signaling component of GABA synthesis [50], flux through GAD67 was set equal to that of GAD65 (and thus 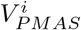):

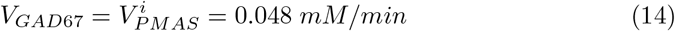

The total flux through the two GAD isoforms was therefore 0.096 mM/min, which is in good agreement with direct estimates of total *V*_*GAD*_ flux in rats [66], after accounting for species differences between human and rodent fluxes.

Consistent with in vitro measurements, 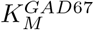 for glutamate was set to 0.13 mM [61–63]. Using Equation 13, with 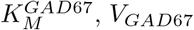 and [*Glu*]_*i*_ known, we determined 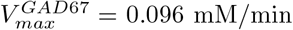. Based on the lack of dependence of GABA concentration on the rate of GABA synthesis when the breakdown enzyme GABA-T is inhibited, we concluded that product inhibition of GAD67 is minimal even within the pharmacological range and therefore did not include an inhibition term in the rate expression [67].

In the reverse pathway of the GABA shunt, 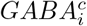 is converted back to glutamate by GABA transaminase (GABA-T, GT), also modeled with Michaelis–Menten kinetics:

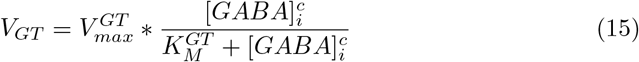

Based on in vitro measurements from prior studies, 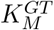 for GABA was set to 0.16 mM [68–70]. Assuming resting-state conditions where *V*_*GAD*67_ = *V*_*GT*_, and 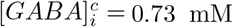, rearranging Equation 15 yielded 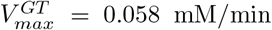. Importantly, GABA-T exhibits lower elasticity to 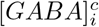 than GAD67 does to [*Glu*]_*i*_. As a result, perturbations in either GAD67 or GABA-T lead to disproportionately large shifts in [*GABA*]^*c*^ due to an amplification effect, as previously observed [67]. This property is a key distinction between GABA and glutamate metabolism, and its downstream consequences are examined in the results section.

### 2.4 Kinetics of the astrocytic reactions of neurotransmitter cycling

Following synaptic release, glutamate and GABA are predominantly taken up by astrocytes for recycling. Within astrocytes, GABA is first converted to glutamate via GABA transaminase (GABA-T), after which the majority of astrocytic glutamate (*Glu*_*a*_) is converted to glutamine (*Gln*_*a*_) and returned to neurons, while a smaller fraction undergoes oxidative metabolism via glutamate dehydrogenase (GDH) [71, 72].

While the reaction catalyzed by GDH is freely reversible, it was assumed to be unidirectional toward glutamate based on prior measurements showing minimal incorporation of ammonia-derived nitrogen into glutamate, even under hyperammonemic conditions [73], suggesting that flux through the reverse reaction is negligible [74]. GDH is strongly regulated through allosteric modulations and it constitutes the link between amino acid metabolism and the TCA cycle [75]. The rate of glutamate oxidation was modeled as a function of [*Glu*]_*a*_ following Hill kinetics:

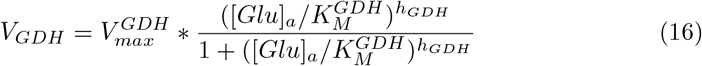

Here, the Hill exponent (*h*_*GDH*_) was set to 4.0 [76], *K*_*m*_ was set to the steady-state value of [*Glu*]_*a*_ = 0.8 mM [45], and 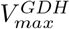 was calibrated to 0.096 mM/min based on in vivo studies reporting that *V*_*GDH*_ equals 20% of the combined neuronal glutamate and GABA release and astrocytic recycle rates under steady-state conditions [36].

Glutamate homeostasis is maintained by replacing oxidized glutamate via anaplerosis. Similarly, glutamine lost through transport-mediated efflux from astrocytes to the extracellular space and from the extracellular space to the blood is also replenished by anaplerosis [72]. Anaplerosis was represented as a composite flux encompassing reactions involved in the synthesis of *α*-ketoglutarate and its subsequent conversion to glutamate through either transamination or reductive amination. We modeled anaplerosis as a constant flux, based on prior studies reporting sustained anaplerotic activity during seizures [77], suggesting that changes in anaplerotic flux occur on timescales longer than those probed in our measurements. The anaplerotic rate was defined as the sum of 20% of the total neuronal glutamate and GABA release rate under resting conditions and the rate of glutamine efflux to plasma (*V*_*out*_) [21]. Under steady-state conditions, *V*_*anapl*_ was 0.06 mM/min, consistent with values reported in the literature [45, 78, 79].

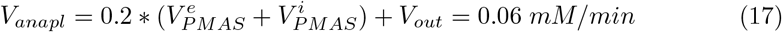

The astrocytic conversion of glutamate to glutamine is mediated by glutamine synthetase (GS). GS activity was described using Michaelis–Menten kinetics as a function of the astrocytic glutamate pool ([*Glu*]_*a*_), with an additional term accounting for product inhibition by glutamine ([*Gln*]_*a*_).

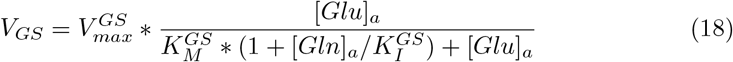

Here, we set 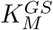 equal to the steady-state value of [*Glu*]_*a*_ = 0.8 mM and 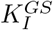 equal to [*Gln*]_*a*_ = 3.8 mM. The value of [*Gln*]_*a*_ was obtained by setting it to 90% of the total [*Gln*], based on intracellular glutamine concentrations in astrocytes and neurons [80] and their relative volume fractions in brain tissue. Future work will be needed to refine the representation of GS kinetics by explicitly accounting for ammonia 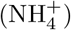, which serves as a substrate for GS, GDH, and PAG within the PMAS and may substantially influence their kinetics [71]. In this work, we assumed 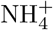 levels remained approximately constant and therefore adopted a simplified representation.

From *Gln*_*a*_, we modeled three outgoing fluxes: two fluxes to the two neuronal compartments, occurring at rates 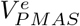 and 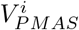, respectively, and a third flux representing glutamine efflux into the blood *V*_*out*_. Therefore, in steady-state:

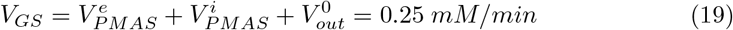

This value is in good agreement with ^13^C MRS data from humans [45]. With this constraint, we determined 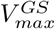 to be 0.75 mM/min based on the maximum increase in total oxidative metabolic rate above the resting awake state that has been measured during extended seizures [29].

The release of glutamine into the blood across the blood brain barrier (BBB) occurs via exchange with branched-chain amino acids and is thought to play an important role in maintaining amino acid homeostasis in the brain [81]. We modeled the outward flux, *V*_*out*_, in two stages, assuming constant levels of substrates exchanged with glutamine.

First, astrocytic glutamine was assumed to maintain a fixed ratio with extracellular fluid levels, reflecting rapid equilibration across the astrocytic membrane via a high-activity transporter that couples the intracellular-to-extracellular glutamine ratio to the Na^+^ gradient, which is maintained relatively constant. This ratio was derived from in vivo microdialysis measurements in resting state, where [*Gln*]_*a*_ = 3.8 mM [45] and [*Gln*]_*ecf*_ = 0.5 mM [82], yielding a ratio of approximately 8:1. Second, glutamine efflux from the extracellular space to plasma across the BBB was described using reversible Michaelis–Menten kinetics with an offset term, *V*_*out*,0_:

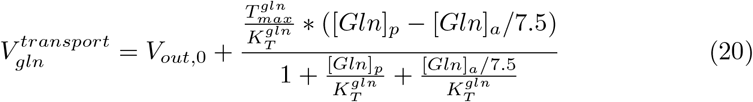

The offset *V*_*out*,0_ was included to account for concentration differences in the counter-transported branched-chain amino acids and was set to 0.012 mM/min based on prior in vivo arteriovenous measurements [83]. Plasma glutamine concentration was taken as [*Gln*]_*p*_ = 0.5 mM [84], while 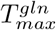 and 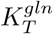 were set to 0.018 mM/min and 0.25 mM, respectively, based on adjusted measurements from rat cortex [85].

### 2.5 Ketone metabolism and its influence on neurotransmitter cycling

In this section, we describe the model components governing the pharmacokinetics of ketones and their effects on neurotransmitter cycling. The specific ketone body considered in both our model and the experimental dataset was D-*β*-hydroxybutyrate (D-*β*HB), which is the primary circulating ketone body under ketogenic conditions. We represented ketone utilization as a two-stage process: the first stage corresponding to uptake across the blood–brain barrier, and the second to turnover through oxidation.

For the first step, we defined the rate of D-*β*HB transport from blood to brain 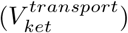 as a function of the plasma ([*Ket*]_*p*_) and brain tissue ([*Ket*]_*b*_) ketone concentrations. To this end, we employed reversible Michaelis–Menten kinetics, following prior formulations developed for lactate transport [86]:

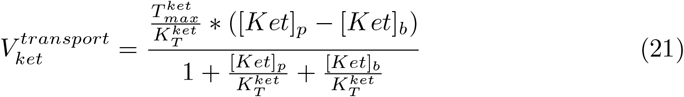

We considered Equation 21 applicable to ketones because both lactate and ketones are transported by the same monocarboxylate transporters (MCTs) and therefore obey analogous kinetics, differing only in their parameter values. While this equation does not explicitly account for competition for the transporters between lactate and ketones, nor for the symport with H^+^, it remains valid as long as the compartmental lactate and H^+^ concentrations remain constant during acute ketosis, which is expected to be the case in the resting awake state. This model component also offers an entry point for future model expansions to study lactate displacement, as well as modeling the impact of lactate on glutamate and GABA levels and related fluxes.

The transport parameters 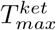 and 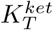 from Equation 21 were determined from two previously reported experiments conducted in humans under strictly non-adapted, acute conditions (SI Table 2). Data obtained from chronic ketogenic states were excluded, as they likely involve additional adaptive mechanisms that are not applicable to the timescales considered in this work. From the two reference studies [7, 34], we extracted two unique triplets of [*Ket*]_*p*_, [*Ket*]_*b*_, and the cerebral metabolic rate of D-*β*HB (*CMR*_*ket*_) values. Assuming steady-state, we set 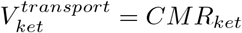 and solved Equation 21 to obtain unique estimates for both parameters: 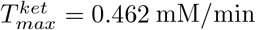 and 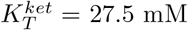.

In the second step, we derived an expression for *CMR*_*ket*_ under the assumption of a linear relationship between *CMR*_*ket*_ and [*Ket*]_*b*_ and a zero intercept (i.e. if [*Ket*]_*b*_ = 0 then *CMR*_*ket*_ = 0). These assumptions are supported by prior in vivo studies in animal models showing that oxidative metabolism has a high affinity for D-*β*HB and does not exhibit saturation within physiological concentration ranges, with saturation first observed at blood levels of ~ 15 mM, well above physiological concentrations [87, 88].

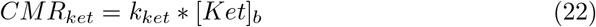

We determined the value of *k*_*ket*_ by fitting Equation 22 to the same two datapoints that we used for calculating the transport parameters [7, 34]. This procedure yielded *k*_*ket*_ = 0.162 min^−1^.

Next, we define the steps for computing *V*_*PMAS*_, the neurotransmitter cycling flux, as a function of *CMR*_*ket*_ via 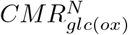. As described above, the PMAS flux is stoichiometrically coupled to the glucose oxidation flux in glutamatergic and GABAergic neurons in order to match neurotransmitter synthesis to neuronal activity and the associated ATP consumption. Prior studies of acute ketosis employing arteriovenous difference measurements [89] and FDG-PET [34] have shown that ketone infusion reduces *CMR*_*glc*(*ox*)_ while maintaining constant cerebral metabolic rate of oxygen. These observations suggest that ketones and glucose compete for the same enzymatic apparatus for oxidation and that ketones partially substitute for cerebral glucose metabolism. We quantified this partial displacement by accounting for the ketogenic contribution to the total oxidative ATP production rate (*V*_*ATP* (*ox*)_), under the assumption that energy production reequilibrates on timescales faster than the kinetics considered in this work. We assumed that under the conditions of acute ketone infusion studied the displacement of glucose oxidation directly impacts the PMAS flux.

First, we determined the value of *V*_*ATP* (*ox*)_ from the baseline *CMR*_*glc*(*ox*)_ (Equation 1) using an ATP yield from glucose 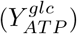 of 32 mol ATP per mol glucose:

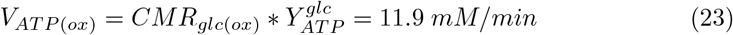

Ketones are metabolized for ATP production exclusively through oxidative metabolism, with an ATP yield 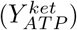 of 21.5 mol per mol D-*β*HB. Accordingly, multiplying *CMR*_*ket*_ by 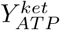 yields the portion of *V*_*ATP* (*ox*)_ displaced by ketone utilization, with glucose oxidation accounting for the remaining portion of total ATP turnover. Combined with Equation 23, this relationship expresses *CMR*_*glc*(*ox*)_ as a function of *CMR*_*ket*_:

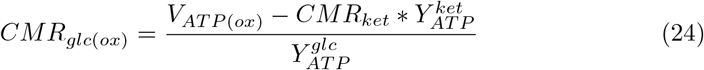

The updated *CMR*_*glc*(*ox*)_ can then be substituted into Equations 2 through 6 to recompute 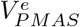 and 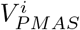 as a function of *CMR*_*ket*_, thereby yielding simulated alterations of neurotransmitter cycling during acute ketosis. The timecourse of *CMR*_*ket*_ can be obtained from measured values of [*Ket*]_*p*_ based on Equation 21. The experimental dataset used in this work for comparisons with simulations includes one such time-series and is described later.

### 2.6 Simulations

The model was implemented in Julia using *Catalyst*.*jl* [90] and *DifferentialEquations*.*jl* [91], with time integration performed using the *Vern7* solver.

To ensure robust simulation results and assess their stability, simulations were repeated across multiple replicates. Input parameters and initial conditions were independently perturbed within a uniform distribution spanning ± 10% around their central values (Table 1 and Table 2). The resulting variability across replicates is represented as error bands in the figures with simulation results.

Each replicate was first simulated for 10,000 minutes with [*Ket*]_*p*_ = 0 to allow the system to reach steady-state prior to ketone administration, after which the ketone bolus was introduced and the simulations were continued for an additional 500 minutes. Replicates resulting in steady-state metabolite concentrations outside the experimentally reported relative uncertainties (total glutamate: ± 9%, GABA: ± 28%) [13] were discarded. This procedure was repeated until a total of 1,000 accepted replicates were obtained.

### 2.7 Experimental dataset

The experimental dataset is from a previously reported 7T ^1^H MRS experiment [12, 13], conducted in a cohort of 63 healthy adults aged 22 to 79 years. Full methodological details are provided in the cited references. Briefly, following overnight fasting, participants ingested (R)-3-hydroxybutyl-(R)-3-hydroxybutyrate ketone monoester at a dose of 395 mg per kg of body weight, diluted in water at a 1:1.6 volume ratio. This compound is rapidly metabolized into D-*β*HB in the body [92]. Five-minute ^1^H MRS scans were acquired immediately before ketone ingestion and again 67 ± 4 (mean ± SD) minutes afterward, with variability arising from practical differences in acquisition timing rather than measurement uncertainty. Brain concentrations of glutamate and GABA were quantified relative to the water signal using LCModel, with correction for cerebrospinal fluid content in the voxel. Percentage changes were then computed relative to the pre-ingestion measurement. Brain levels of glutamate decreased by 10.6% ± 2.4% (mean ± 95% CI), while GABA levels showed a larger decrease of 36.9% ± 8.2%.

To obtain model outputs compatible with the experimental data at *t* = 67 min, a time series of blood ketone levels was required as input for the model. As with the brain measurements, however, the experimental dataset included only a single blood measurement at *t* = 75 min rather than a full time series. To acquire time-series, we adopted a temporal curve from a comparable study that measured blood ketone dynamics following a similar dose (360 mg per kg of body weight) in healthy adults [93]. The closest reported value in this study at *t* = 60 min (4.08 mM) was in good agreement with the corresponding blood value from the MRS dataset at *t* = 75 min (4.16 mM ± 0.133 mM; mean ± 95% CI), and therefore we applied no additional scaling. The resulting curves were also consistent with prior reports describing comparable blood ketone dynamics under similar administration paradigms [94, 95]. Finally, we fitted the published data with a double exponential function to generate a continuous input series as input data for the model (Figure 2A) where *t* represents the elapsed time in minutes:

**Fig. 2.**
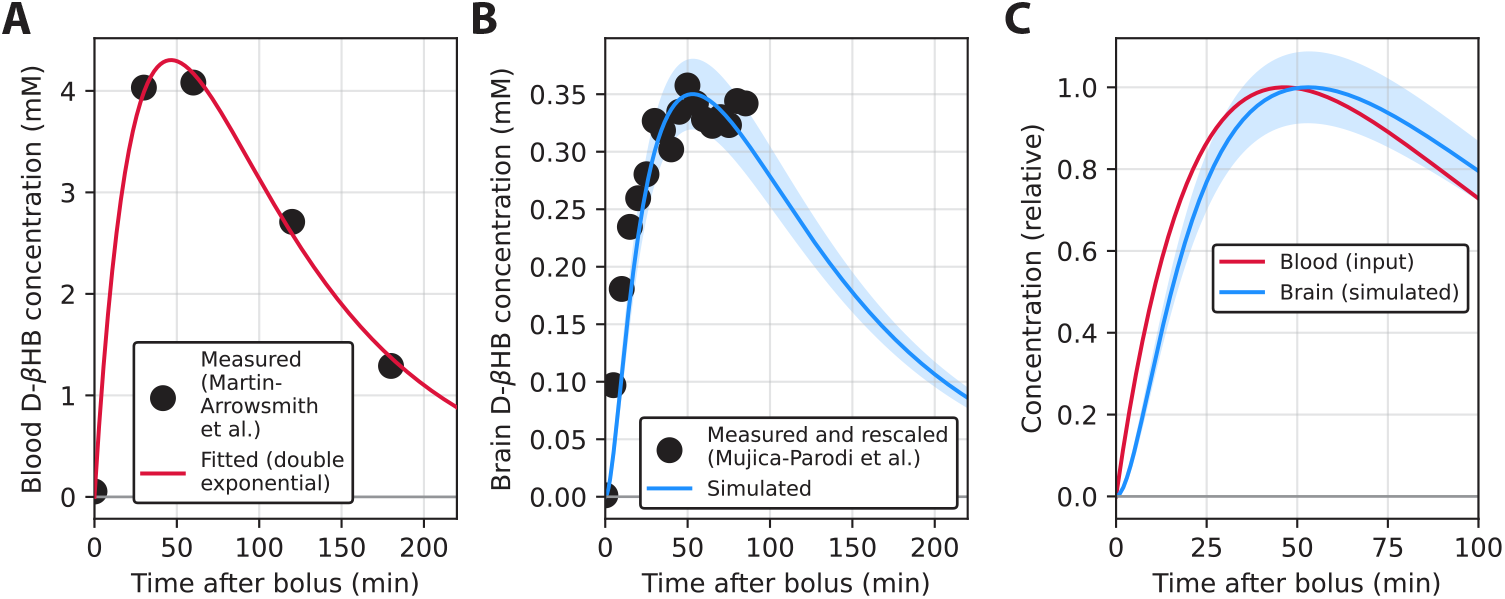
Measured and simulated time courses of brain ketone metabolism. **A**: Blood D-*β*HB levels (black dots) following acute ketone administration, obtained from the literature [93], fitted with a double exponential function (red). **B**: Simulated brain D-*β*HB concentrations (blue), generated by the model using the blood input from panel A. For reference, brain D-*β*HB concentrations measured with MRS (black dots) are shown after rescaling to match the scale of the simulated values, since the MRS data were collected in arbitrary units [96]. The error band for the simulated curve represents the standard deviation obtained by varying all model parameters by ± 10% around their central values. **C**: Overlay of the fitted blood D-*β*HB curve and simulated brain D-*β*HB concentrations from panels **A** and **B**, normalized to the maximum of each curve, highlighting the short ~ 5-minute lag between the two.

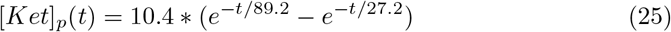

MRS measurements of brain ketone levels from a prior study were also used to validate the intermediate model outputs of brain ketone concentration time series [96]. These data were obtained from an MRS experiment independent of the main experimental dataset but conducted under the same ketone administration paradigm. Because these measurements were reported in arbitrary units, they served as a reference for relative dynamics rather than absolute concentrations.

### 2.8 Metabolic control analysis

Metabolic control analysis (MCA) is an approach to quantify the influence of parameters on fluxes and metabolite concentrations [97]. Its main utility is in identifying the most influential parameters that require careful description and parameterization (and can serve as entry points for interventions), while also revealing components that can be simplified or omitted due to limited influence. MCA is analogous to sensitivity analysis, with the important distinction that it quantifies parameter influence on the steady-state, after the system’s elements have interacted and converged. Although control coefficients correspond to individual parameters, metabolic control is fundamentally a system property. Flux control coefficients describe the dependence of steady-state fluxes on parameters, whereas concentration control coefficients describe the dependence of steady-state metabolite concentrations. Conventionally, control coefficients are defined with respect to enzyme concentrations, but analogous measures can be computed for other parameters, such as Michaelis–Menten constants, in which case they are termed response coefficients.

While flux control in our model was trivially confined to the PMAS and altered only by ketones, concentration control was expected to be more broadly distributed across reactions. We quantified concentration control coefficients using the expression described in [37], where the coefficient 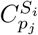 denotes the influence of parameter *p*_*j*_ on the steady-state concentration of metabolite *S*_*i*_.

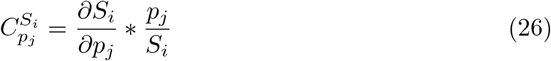

We solved this equation numerically using a central finite difference approximation.

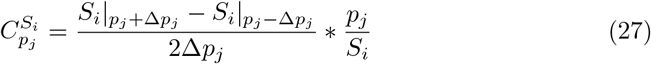

We chose *ϵ*, the relative perturbation, where *ϵ* = Δ*p*_*j*_*/p*_*j*_, to be 1%. After substitution, the numerical solution simplifies to:

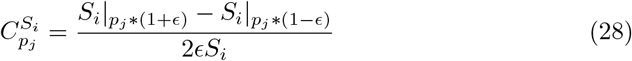

## 3 Results

### 3.1 Predicting the rate of brain ketone utilization after an acute challenge

Our first step was to establish the model component responsible for simulating ketone kinetics in the brain, which then served as input to the full neurotransmitter cycling model. To this end, we first simulated the brain tissue concentration of D-*β*HB in response to acute ketone administration, using the time-series of blood D-*β*HB concentration as input (Figure 2A). The simulated brain dynamics closely matched the rescaled values of the experimental dataset of brain D-*β*HB levels [96] (rescaled due to arbitrary units), particularly the rapid increase immediately after ketone administration and the timing of the peak (Figure 2B). Notably, both experimental and simulated data showed that brain concentrations closely tracked blood concentrations, lagging by only approximately 5 minutes (Figure 2C). This observation is consistent with prior studies showing that brain D-*β*HB concentrations are low relative to transport, which implies high turnover [87]. The simulated brain ketone concentration reached its peak 53 minutes after the ketone bolus.

### 3.2 Model predictions of glutamate and GABA agree with experimental data, supporting the PMAS-centered hypothesis

Next, we proceeded to our full model of neurotransmitter cycling to simulate dynamic changes in glutamate and GABA levels following an acute D-*β*HB bolus for comparison with the experimental dataset. Figure 3 shows the predicted changes from initial steady-state fluxes and concentrations, generally peaking 50–200 minutes after ketone administration. As blood ketone levels dropped, all of the simulated variables eventually reconverged to baseline, although only after several hours of simulated time. Longer time courses for all metabolites are provided in SI Figure 1.

**Fig. 3.**
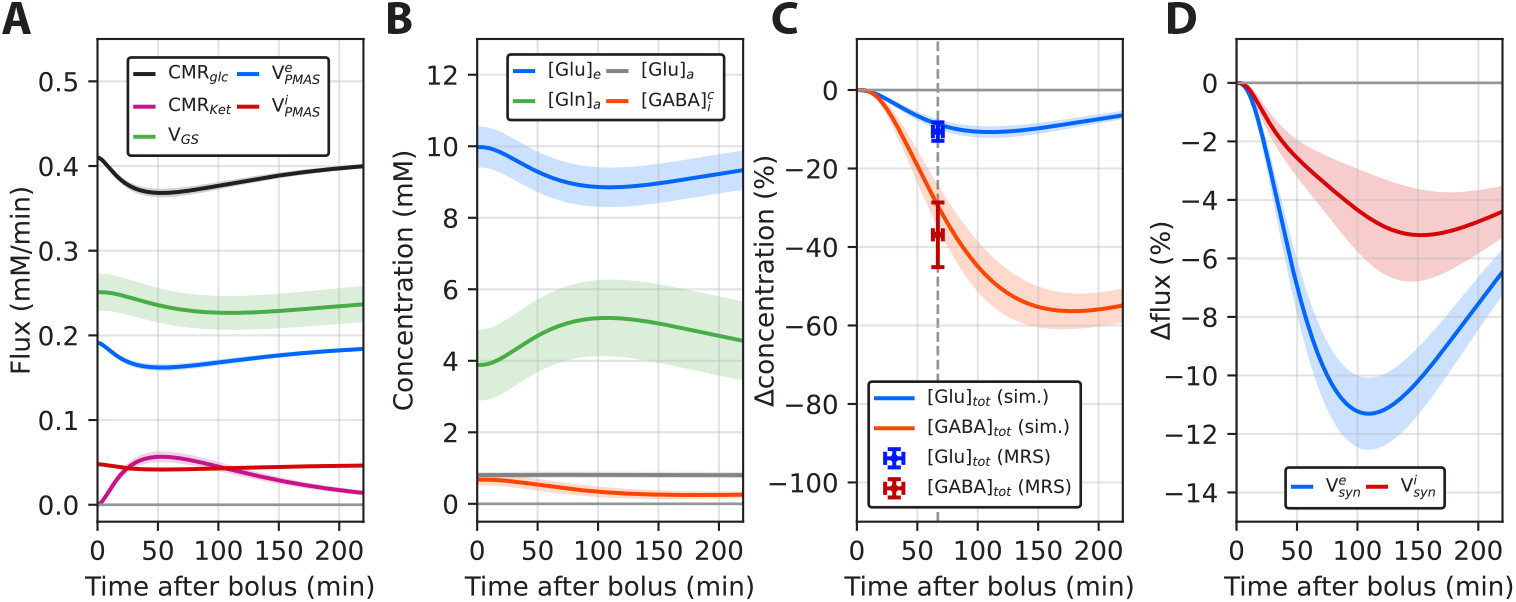
Simulated time courses of neurotransmitter cycling fluxes and concentrations following an acute ketone bolus, compared with MRS measurements. **A**: Time courses of selected fluxes after a D-*β*HB bolus, highlighting the varying magnitudes of change and the distinct temporal delays across reactions within the cycling pathway. Error bands for the simulations represent the standard deviation obtained by varying parameters by *±*10% around their central values. Shown are the cerebral metabolic rate of glucose (*CMR*_*glc*_) and ketones (*CMR*_*ket*_), the rate of glutamine synthetase (*V*_*GS*_), and fluxes through the pseudo-malate-aspartate shuttle in glutamatergic and GABAergic neurons (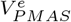 and 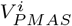). **B**: Time courses of absolute concentrations of selected compartmental pools of glutamate, glutamine, and GABA. The small pools of [*Glu*]_*i*_, [*Gln*]_*e*_, [*Gln*]_*i*_, and 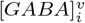 are shown in SI Figure 1. **C**: Time courses of relative changes in the total (sum of all compartments) glutamate (blue) and GABA concentrations (red), compared with MRS measurements from the experimental dataset. Changes are expressed as percentages to allow comparison with the MRS results, which were originally reported in arbitrary units. Vertical error bars on the measurement points show 95% confidence intervals across subjects. Horizontal error bars reflect the standard deviation of sampling times, which varied across participants due to practical timing differences. **D**: Simulated time courses of relative changes in synaptic glutamate (blue) and GABA (red) fluxes.

Based on the model, the predicted decline in *CMR*_*glc*_ was concurrent with the rise in *CMR*_*ket*_ (Figure 3A), consistent with prior observations that ketones partially displace glucose metabolic flux [34]. *CMR*_*glc*_ decreased from 0.41 to 0.37 mM/min (−10%) on average, reaching its minimum at 53 minutes, while *CMR*_*ket*_ peaked at0.056 mM/min. Fluxes through 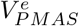 and 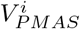 also decreased, due to their coupling with 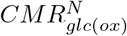, from 0.19 to 0.16 mM/min and from 0.048 to 0.042 mM/min, respectively. These simulated changes were followed by a reduction in flux through GS, decreasing from 0.25 to 0.23 mM/min and reaching a minimum at 110 minutes. In contrast, GABAergic reactions involving GAD65, GAD67, and GABA-T responded more slowly to ketones in the simulations, with mean minima occurring between 151 and 176 minutes. Notably, fluxes through GAD67 and GABA-T, which together represent an inner GABA cycle, decreased substantially from 0.048 to 0.034 mM/min at their minima, whereas flux through GAD65 remained relatively stable, decreasing only from 0.048 to 0.045 mM/min.

In line with the predicted flux alterations, the metabolic concentration pools also exhibited substantial changes in response to ketones. The glutamatergic neuronal glutamate pool ([*Glu*]_*e*_), which supports excitatory synaptic function, underwent a marked decrease, reaching a minimum 109 minutes after the ketone bolus (10.0 to 8.85 mM, −11%) (Figure 3B). The astrocytic glutamate pool ([*Glu*]_*a*_), which maintains equilibrium with the neuromodulatory extracellular glutamate pool [98], remained relatively stable (relative changes within 1%), while the astrocytic glutamine pool ([*Gln*]_*a*_) increased by 34% on average, primarily due to the delayed reduction in flux through GS relative to PMAS. Meanwhile, the cytosolic GABA pool 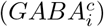, which constitutes the majority of the measurable GABA and is in equilibrium with neuromodulatory extracellular GABA, declined from 0.73 to 0.25 mM (−66%). This reduction was not only substantially larger than that observed for glutamate, but also occurred on a slower timescale, reaching a minimum at 183 minutes.

The observed compartmental changes were also reflected in the total metabolite pools and, importantly, showed good agreement between model predictions and experimental results (Figure 3C). For comparability with the MRS data, compartmental pools were merged into total metabolite concentrations: [*Glu*]_*tot*_ was calculated as the sum of [*Glu*]_*e*_, [*Glu*]_*i*_, and [*Glu*]_*a*_, whereas [*GABA*]_*tot*_ was calculated as the sum of 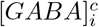 and 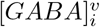. The predicted reduction in total brain glutamate concentration at 67 minutes post ketone bolus was 9%, within the 95% CI of the observed decrease (10.6% *±* 2.4%). GABA exhibited a substantially larger decline, with the model predicting a 30% decrease at 67 minutes, also within the 95% CI of the experimental estimate (−36.9% ± 8.2%). The agreement between the model predictions and experimental observations provides quantitative support for our hypothesis that acute ketosis modulates neurotransmitter cycling through reductions in *CMR*_*glc*(*ox*)_ and PMAS activity.

Although the total GABA pool showed a substantially larger relative reduction than the glutamate pool, this did not translate to the synaptic fluxes, where the glutamate-driven fluxes underwent the greater decrease (Figure 3D). This divergence arose from the bifurcation of the GABA pools: synaptic GABA release is supplied by the vesicular GABA pool, which is distinct from the cytosolic pool and remained comparatively stable, exhibiting only a 5% decrease at 152 minutes (SI Figure 1). These observations suggest that although the measurable total GABA concentration may decline more strongly than glutamate during ketosis, the synaptic GABA flux actually remains comparatively robust and undergoes a smaller reduction than the synaptic glutamate flux.

### 3.3 Metabolic control analysis identifies key control points for regulating glutamate and GABA levels

With the model predictions established to be consistent with experimental data, we next asked which parameters exert the strongest influence on neurotransmitter homeostasis. While the control of neurotransmitter cycling flux was tied to 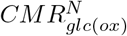 through the PMAS (a core assumption of the model introduced earlier, see Equation 5 and Equation 6), we expected concentration control to be more distributed across the system. To assess how specific parameters influence metabolite and neurotransmitter concentrations, we carried out metabolic control analysis in the control state, with no ketones present in the system (Figure 4). We focused on key parameters, including *V*_*max*_ terms associated with enzymatic activity, which are potential entry points to genetic and therapeutic regulation, and *K*_*M*_ parameters reflecting intrinsic kinetic properties of the enzymes. We also introduced a weighting parameter for *V*_*PMAS*_, used only in this analysis, to assess the influence of ketosis, which was hypothesized to affect neurotransmitter cycling through PMAS, and to evaluate our hypothesis that PMAS is the primary control point for glutamate and GABA concentrations under these conditions.

**Fig. 4.**
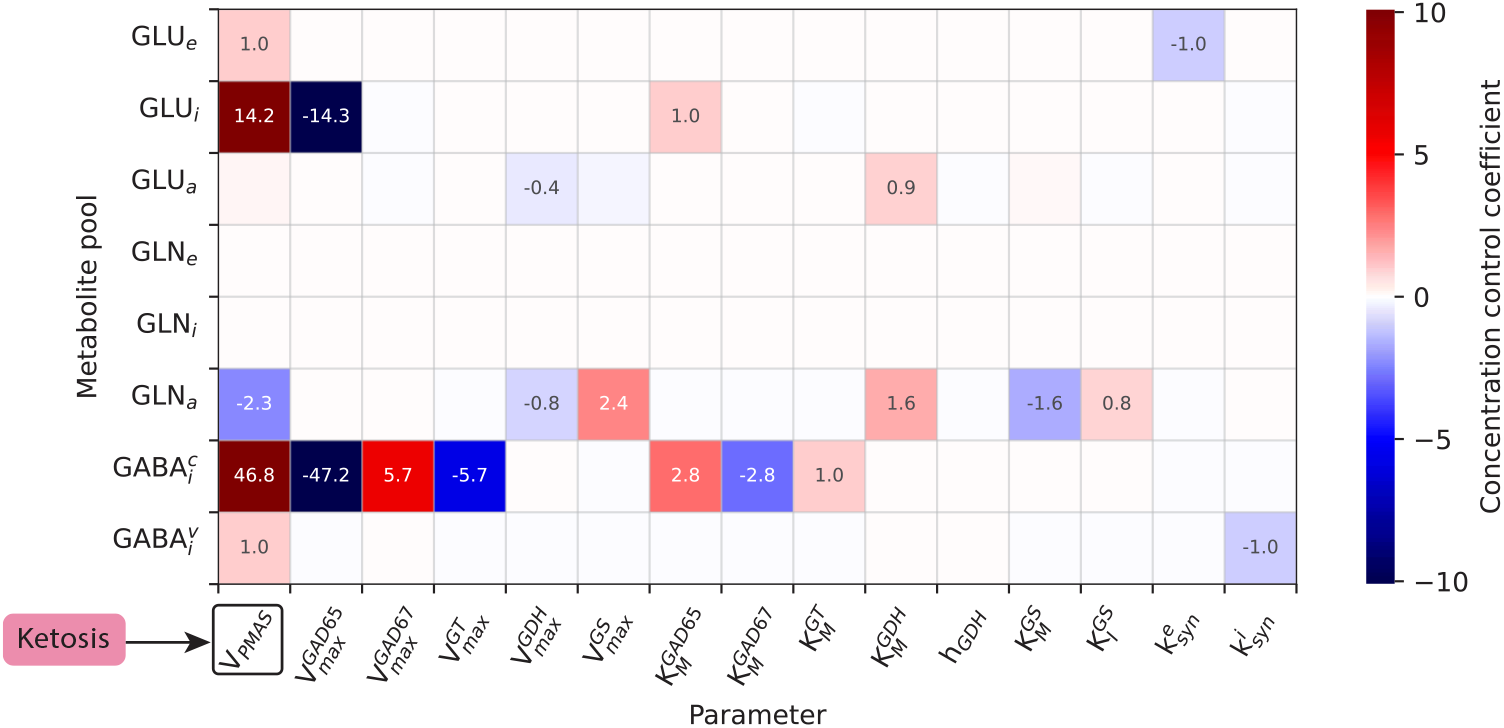
Metabolic control analysis of metabolite pools in neurotransmitter cycling. The heatmap indicates the degree of metabolic control on concentration exerted by a parameter (horizontal axis) on each metabolite (vertical axis). A value of +1 indicates that a +1% change in the parameter value yields a +1% change in the metabolite steady-state concentration, while values around zero indicate the absence of control. Only a subset of parameters is shown here, including *V*_*max*_ values, which primarily reflect enzyme concentration or maximum activity, and kinetic parameters determining enzyme elasticity (e.g., *K*_*M*_). Here, *V*_*PMAS*_ represents a weighting parameter for the PMAS rate, intended to highlight the modeled control exerted by ketosis on the metabolite pools (represented schematically by an arrow from “Ketosis” to *V*_*PMAS*_).

The computed concentration control coefficients (Figure 4) indicated that the rate of PMAS (*V*_*PMAS*_), through which ketosis was hypothesized to act, exerted significant control over multiple metabolite concentrations, particularly 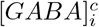 and [*Glu*]_*i*_ in the GABAergic neuron, and to a lesser extent [*Glu*]_*e*_, 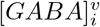, and the astrocytic glutamine pool [*Gln*]_*a*_. Beyond the shared regulation of glutamate and GABA, the model identified parameters that selectively regulate GABA, particularly [*GABA*]^*c*^, with minimal influence on glutamate. These included the GABAergic neuron–specific enzymes GAD65, GAD67, and GABA-T. Because GAD65 activity is expected to increase in parallel with *V*_*PMAS*_ during activation, this pattern was expected. In astrocytes, GDH influenced both [*Glu*]_*a*_ and [*Gln*]_*a*_, whereas the activity of GS controlled its product [*Gln*]_*a*_ but not its substrate [*Glu*]_*a*_.

To assess the robustness of these observations, we repeated the analysis under the ketosis state ([*Ket*]_*p*_ fixed at 4.08 mM) and obtained an equivalent, though attenuated, distribution of control (SI Figure 2), indicating that the identified control points are preserved across conditions. Overall, these results provide further support for our hypothesis, and they identify multiple enzyme steps that can selectively impact compartmental glutamate and GABA levels, establishing them as potential metabolic and pharmacological intervention targets.

## 4 Discussion

In the brain, glutamate and GABA are regulated by neuron- and astrocyte-mediated biochemical reactions that are tightly coupled to core metabolic pathways [21]. This linkage offers an entry point for metabolic interventions aimed at modulating neurotransmitter levels. Ketones, as alternative energy substrates for the brain, present a promising avenue for such interventions [99]. A recent in vivo MRS study has shown that *β*-hydroxybutyrate reduces brain levels of glutamate and GABA within an hour of administration, supporting the therapeutic potential of ketones [12, 13]. However, the underlying mechanism remained unknown and is difficult to probe with existing experimental paradigms. In this work, we present a computational model as a way to bridge this gap, demonstrating how modeling can be used to test potential mechanistic explanations for these observed effects and, more broadly, to provide insights into the metabolic regulation of neurotransmitter homeostasis.

At the core of our model was a kinetic description of neurotransmitter cycling including the three major compartments (glutamatergic neurons, GABAergic neurons, and glia), as well as transport fluxes to the extracellular fluid and the plasma across the blood brain barrier. Given the multitude of reactions involved in this pathway [21, 65], incorporating all of them with accurate parameters is infeasible due to limitations in available experimental data. To address this, we applied the principle of metabolic control analysis, which posits that the majority of control over a metabolic pathway is typically exerted by a subset of reactions [97]. Accordingly, only these reactions require a detailed kinetic representation to simulate pathway behavior accurately. This notion is also key to scaling up computational models, as it reduces the number of required equations and makes scaling tractable. We applied the same principles to formulate the other major component of our model: a pharmacokinetic description of ketone metabolism, including the partial displacement of glucose as an energy substrate [7, 34, 86]. We then connected this component to neurotransmitter cycling by linking oxidative glucose flux, modulated by ketone metabolism, to glutamate synthesis via the pseudo-malate-aspartate shuttle [32]. The complete model was subsequently parameterized using values from prior experimental studies, primarily from in vivo measurements. Importantly, to avoid circularity, the MRS dataset used to test for consistency with model predictions were not involved in model construction.

By matching the dosing and timescale of the reference experiment to measured blood ketone levels, our model yielded results in good agreement with the observed 11% decrease in glutamate and 37% reduction in GABA [13]. Therefore, our results provide quantitative support for the proposed mechanism in which ketones partially displace glucose oxidation, leading via the pseudo-malate-aspartate shuttle to reduced neurotransmitter cycling and, consequently, lower steady-state concentrations of glutamate and GABA. This interpretation was also consistent with previous work implicating the malate-aspartate shuttle in mediating ketone-induced modulation of glutamate metabolism [100].

Prior experimental evidence suggests there may be limits to the extent to which ketones can displace glucose oxidation. A study simultaneously measured glutamate and GABA neurotransmitter fluxes at different levels of brain activity in rat cerebral cortex under conditions ranging from no ketone displacement to maximal ketone-induced displacement of glucose oxidation [87]. They found that ketones can replace glucose oxidation in supporting all non-signaling components of brain ATP consumption. However, the majority of ATP required to support signaling-related energy metabolism remained dependent on glucose oxidation. Based on this study, together with previous studies examining the maximum displacement of brain glucose metabolism by ketones in awake humans [101], the magnitude of PMAS inhibition predicted in the present study may represent an upper bound, potentially protecting glutamate and GABA concentrations from excessive depletion. Further experimental studies will be needed to test this possibility.

The comparatively larger reduction in GABA can be explained by an amplification effect arising from the different elasticities of GAD67 and GABA-T [67], also consistent with the metabolic control analysis results that highlighted selective control of cytosolic GABA by GAD67 and GABA-T. These enzymes catalyze key reactions within a cycle internal to GABAergic neurons, thereby coupling their fluxes. A reduced supply of glutamate to GABAergic neurons decreases flux through GAD67. However, due to the low sensitivity of GABA-T to its substrate GABA, the system reaches a new equilibrium only at a substantially lower cytosolic GABA concentration, resulting in an amplified depletion of this pool. Consistent with this interpretation, similar amplification effects have been reported in studies of selective GABA-T inhibitors, such as vigabatrin [102, 103].

The metabolic control analysis also indicated that GAD65 exhibited a particularly high control coefficient for cytosolic GABA, which was unexpected given its relatively limited role in modulating the measurable GABA concentration as shown in knock out models [104], and the stability of GABA levels in vivo across large changes in brain activity and GAD65 flux [29, 50]. We propose that the likely explanation for the high sensitivity of cytosolic GABA concentration to GAD65 arises because acute ketone elevation selectively inhibits the 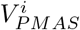, thereby creating a mismatch between glutamate synthesis via 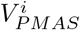 and the flux of GABA leaving the cell through GAD65 and vesicular release. This mismatch leads to depletion of cytosolic GABA until intracellular glutamate reaches a level sufficiently low to reduce the GAD65 flux to match 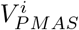. The resulting reduction in intracellular glutamate leads to a decrease in GAD67 flux, which in turn produces the previously described amplification effect on 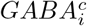 due to the low elasticity of GABA-T. In contrast, we propose that during activity-driven physiological changes in GABA neurotransmitter cycling and metabolism, the fluxes through PMAS, GAD65, and GABAergic vesicle release are proportionally activated and deactivated, resulting in the minimal impact of such changes on GABA and glutamate concentrations [29, 50]. Direct evidence for proportional activation of GAD has been provided by studies showing activity dependent activation of GAD65 via phosphorylation [105].

The 37% reduction in GABA observed in the experimental dataset was profound. Although such a large effect is unlikely to persist at longer timescales as adaptation mechanisms eventually counteract it, changes in GABA of similar magnitudes have previously been observed on acute timescales, including a 35% increase in humans following selective serotonin reuptake inhibitor administration [106], as well as an approximately threefold increase in GABA both rat models [67] and humans [103] in response to vigabatrin. Thus, whether there is sufficient time for adaptation to occur may explain why some studies have reported seemingly contradictory findings, such as increases in GABA during chronic ketosis [10, 11, 107]. When such adaptations are accounted for, we expect the simulated GABA levels to return to baseline more rapidly than predicted by simulations of metabolic processes alone. Longer-term adaptation can be directly incorporated into the model by adjusting the total activities and kinetic constants for the enzymes included. Based on the control analysis, which pinpointed key concentration controlling reactions, we anticipate that a reduced subset of enzymatic reactions will be sufficient, lowering the number of studies required to parameterize the model.

Our model predicted a clear difference between neuromodulatory and synaptic neurotransmitter pools in their response to acute ketosis. While cytosolic GABA, which has a neuromodulatory role but minimal impact on the vesicular pool GABA concentration, showed a pronounced decrease, the vesicular GABA pool that directly interfaces with inhibitory synaptic function was predicted to remain stable as a consequence of the kinetics of GAD65 and potentially the GABA vesicular transporter [48, 49]. This behavior contrasts with glutamate, for which the vesicular glutamate pool in excitatory neurons, which supports excitatory synaptic signaling, was predicted to undergo a substantial reduction. These results highlight the utility of differentiating between synaptic and neuromodulatory pools of glutamate and GABA, as our model suggests strongly divergent responses to perturbations in neurotransmitter cycling. In the case of GABA, the reduction was specific to the neuromodulatory pool, which constitutes the majority of total GABA and therefore dominates the MRS signal, whereas the synaptic vesicular GABA pool remained minimally affected. This distinction may help explain the apparent contradiction between experimental observations of reduced GABA measured by MRS during ketosis [13], which could be interpreted as reduced inhibitory function, and the anticonvulsant effects of ketones, which are generally associated with reduced excitatory signaling [1]. The reduction in synaptic glutamate concentration may compensate for the drop in neuromodulatory cellular GABA during seizures, when the large increase in excitatory glutamatergic signaling transiently elevates extracellular and astrocytic glutamate pools. If, unlike glutamate, the vesicular GABA pool remains robust to ketones, the net effect, even in the presence of reduced neuromodulatory GABA, may be a reduction in excitatory signaling rather than impaired inhibition. Future work will be needed to further clarify the distinct functional roles of synaptic and neuromodulatory pools of glutamate and GABA and to better link metabolic observations with functional outcomes.

While computational models can help narrow the range of viable hypotheses, experimental confirmation remains essential. We selected our experimental dataset because, from a mechanistic standpoint, it was the most suitable dataset available for capturing the direct effects of ketones on neurotransmitters in vivo with the least confounding influence [13]. For example, the study was carried out in healthy adults, thereby avoiding the potential confounding of altered metabolism in patient populations. The short timescale investigated ( ~ 1 hour) also minimized the influence of metabolic and genetic adaptation mechanisms, which prior work has shown to depend strongly on the duration of the intervention [108–110]. Indeed, experimental studies have reported higher brain ketone concentrations under adapted conditions [88] compared to acute states [7], likely reflecting upregulation of ketone transport with prolonged fasting. Conversely, other adaptation mechanisms may counteract the effects on neurotransmitters and dampen their magnitude.

Although the simulation results showed good agreement with experimental data, alternative possibilities remain for how ketones exert modulatory effects on neurotransmitter cycling on acute timescales. It is also possible that several mechanisms contribute simultaneously, motivating future expansions of the model. One set of such mechanisms relates to shifts in the balance between glutamate and *α*-ketoglutarate in the presence of ketones [11, 111, 112]. Given that ketones are not anaplerotic, unlike glucose, a transition to ketogenic states may also reduce anaplerotic flux, thereby limiting the supply of neurotransmitter pools and ultimately contributing to reduced glutamate and GABA levels [12, 113]. Other studies have reported effects of ketones on vesicular signaling, which could further influence neurotransmitter cycling rates [114, 115]. Finally, there is increasing evidence highlighting the role of brain glycogen as an energy source on fast timescales [29, 116]. By partially displacing glucose, the substrate for glycogen synthesis, ketones may modulate glycogen metabolism, potentially exerting distinct effects across different neuronal populations. In particular, fast-spiking inhibitory interneurons, which are known to have high metabolic demand and rapid neurotransmitter cycling [117], may be affected to a greater extent by this mechanism, consistent with the stronger effects observed for GABA.

To our knowledge, this is the first computational model capable of simultaneously evaluating competition between energy substrates in the brain and their impact on neurotransmitter cycling, as well as on intracellular, vesicular, and extracellular concentrations of glutamate and GABA. This integration enables *in silico* evaluation of ketogenic interventions in disease contexts where neurotransmitter imbalance plays an important role, such as schizophrenia [118, 119] and bipolar disorder [120]. With a clearer understanding of how ketones exert these effects, ketogenic therapies may be paired more effectively with treatments targeting complementary mechanisms to improve clinical outcomes. Once established as treatments with known mechanisms, ketogenic interventions could reduce reliance on trial-and-error prescribing and allow for lower doses, which is a meaningful advantage given the often substantial side effects of psychiatric medications [121, 122]. A major utility of our quantitative model is its ability to evaluate hypotheses involving specific dosing and timing of acute ketogenic interventions. Using our tool, one can ask: for a given desired effect, what plasma ketone concentration must be achieved? Altogether, these capabilities offer a foundation for positioning ketogenic therapies more effectively for clinical translation.

On a broader scale, our model could serve as a foundation for simulating neurotransmitter homeostasis across a variety of contexts. There is a growing need for such frameworks, as the rapidly increasing volume of experimental findings is becoming difficult to fully leverage using conventional approaches alone. Computational models, in contrast, allow for the integration of diverse data sources and enable the testing of hypotheses that rely on multiple lines of evidence simultaneously. A key utility of integrating metabolic control analysis is its ability to pinpoint new drug targets by identifying enzymes with the strongest influence over a pathway, thereby revealing the most effective points of intervention. Biomarker discovery represents another major application, particularly the identification of early-stage indicators of disease. In brain aging, for instance, emerging evidence suggests that pathological changes begin well before clinical symptoms appear [5, 123, 124], highlighting the need for early biomarkers that enable timely and effective interventions. Such early-stage neurodiagnostics are likely to be detectable with functional neuroimaging modalities, including functional magnetic resonance imaging (fMRI) and electroencephalography (EEG), as neuronal function is expected to exhibit sensitivity to pathology earlier than structural changes [125, 126]. Consistent with this trend, research increasingly employs computational models of neuronal dynamics and brain circuit behavior [127]. Because neurotransmitters form a critical link between metabolism and neuronal function, our model offers a natural interface for integration with models of brain activity, enabling the propagation of metabolic changes through to neuroimaging signals and even behavioral outcomes.

To assist in using the model presented here for further studies of the neuromodulatory mechanisms of ketosis, as well as other interventions targeted at metabolism, the model presented here is embedded within Neuroblox (https://www.neuroblox.ai/) [128, 129], a computational neuroscience platform designed for data-driven brain modeling. Neuroblox includes a wide range of neuronal function models and offers a user-friendly graphical interface, allowing researchers to interact with and extend our model for diverse applications. The implementation exposes all model parameters to user modification and supports user-defined input time courses.

In conclusion, we presented a novel computational modeling framework integrating dynamic representations of neuronal metabolism and neurotransmitter cycling. Using this framework, we tested a mechanistic hypothesis for how ketones influence the neurotransmitters glutamate and GABA. Beyond the specific findings presented here, our model may also serve as a foundation for future integrative computational approaches aimed at extracting mechanistic insights from the rapidly growing body of experimental data in this field.

## 5 Data and Code

The model presented in this work is part of the Neuroblox computational neuroscience platform (code: https://neuroblox.ai/code; documentation: https://neuroblox.ai/docs). An easy-to-use GUI, tutorials, and a standalone implementation for reproducing the results in this work are available at https://neuroblox.ai/pubs.

## 6 Acknowledgments

The research presented here was funded by the Baszucki Foundation (supporting LRMP). LRMP are HHS are co-founders of Neuroblox Inc. EMR is a consultant for Aletheia and is an unpaid member on the advisory board of BrainSpec. We thank Kevin Behar for helpful discussions regarding the parameters used in the modeling. We also thank Alex Driussi for the scientific illustration presented in Figure 1.

## References

[1] Neal, E. G. et al. The ketogenic diet for the treatment of childhood epilepsy: a randomised controlled trial. The Lancet Neurology 7, 500–506 (2008).

[2] Kraeuter, A.-K., Phillips, R. & Sarnyai, Z. Ketogenic therapy in neurodegenerative and psychiatric disorders: From mice to men. Progress in Neuro-Psychopharmacology and Biological Psychiatry 101, 109913 (2020).

[3] Fortier, M. et al. A ketogenic drink improves cognition in mild cognitive impairment: Results of a 6-month rct. Alzheimer’s & Dementia 17, 543–552 (2021).

[4] Murray, A. J. et al. Novel ketone diet enhances physical and cognitive performance. The FASEB Journal 30, 4021 (2016).

[5] Antal, B. B. et al. Brain aging shows nonlinear transitions, suggesting a midlife “critical window” for metabolic intervention. Proceedings of the National Academy of Sciences 122, e2416433122 (2025).

[6] Croteau, E. et al. A cross-sectional comparison of brain glucose and ketone metabolism in cognitively healthy older adults, mild cognitive impairment and early alzheimer’s disease. Experimental gerontology 107, 18–26 (2018).

[7] Pan, J. W. et al. [2, 4-13c2]-β-hydroxybutyrate metabolism in human brain. Journal of Cerebral Blood Flow & Metabolism 22, 890–898 (2002).

[8] Kolen, B. et al. Vesicular glutamate transporters are h+-anion exchangers that operate at variable stoichiometry. Nature Communications 14, 2723 (2023).

[9] Roth, F. C. & Draguhn, A. Gaba metabolism and transport: effects on synaptic efficacy. Neural plasticity 2012, 805830 (2012).

[10] Yudkoff, M. et al. The ketogenic diet and brain metabolism of amino acids: relationship to the anticonvulsant effect. Annu. Rev. Nutr. 27, 415–430 (2007).

[11] Qiao, Y.-N. et al. Ketogenic diet-produced β-hydroxybutyric acid accumulates brain gaba and increases gaba/glutamate ratio to inhibit epilepsy. Cell Discovery 10, 17 (2024).

[12] Hone-Blanchet, A. et al. Acute administration of ketone beta-hydroxybutyrate downregulates 7t proton magnetic resonance spectroscopy-derived levels of anterior and posterior cingulate gaba and glutamate in healthy adults. Neuropsychopharmacology 48, 797–805 (2023).

[13] van Nieuwenhuizen, H. et al. Ketosis elevates antioxidants and markers of energy metabolism: A 1h mr spectroscopy study. Biological Psychiatry: Cognitive Neuroscience and Neuroimaging (2025).

[14] Mosconi, L., Pupi, A. & De Leon, M. J. Brain glucose hypometabolism and oxidative stress in preclinical alzheimer’s disease. Annals of the New York Academy of Sciences 1147, 180–195 (2008).

[15] Wang, W., Zhao, F., Ma, X., Perry, G. & Zhu, X. Mitochondria dysfunction in the pathogenesis of alzheimer’s disease: recent advances. Molecular neurodegeneration 15, 1–22 (2020).

[16] Austin, B. P. et al. Effects of hypoperfusion in alzheimer’s disease. Journal of Alzheimer’s Disease 26, 123–133 (2011).

[17] Videbech, P. Pet measurements of brain glucose metabolism and blood flow in major depressive disorder: a critical review. Acta Psychiatrica Scandinavica 101, 11–20 (2000).

[18] Townsend, L. et al. Brain glucose metabolism in schizophrenia: a systematic review and meta-analysis of 18fdg-pet studies in schizophrenia. Psychological Medicine 53, 4880–4897 (2023).

[19] Chesebro, A. G., Antal, B. B., Weistuch, C. & Mujica-Parodi, L. R. Challenges and frontiers in computational metabolic psychiatry. Biological Psychiatry: Cognitive Neuroscience and Neuroimaging (2024).

[20] Patel, A. B. et al. The contribution of gaba to glutamate/glutamine cycling and energy metabolism in the rat cortex in vivo. Proceedings of the National Academy of Sciences 102, 5588–5593 (2005).

[21] Andersen, J. V. et al. Glutamate metabolism and recycling at the excitatory synapse in health and neurodegeneration. Neuropharmacology 196, 108719 (2021).

[22] Harris, J. J., Jolivet, R. & Attwell, D. Synaptic energy use and supply. Neuron 75, 762–777 (2012).

[23] Yu, Y., Herman, P., Rothman, D. L., Agarwal, D. & Hyder, F. Evaluating the gray and white matter energy budgets of human brain function. Journal of Cerebral Blood Flow & Metabolism 38, 1339–1353 (2018).

[24] Calvetti, D. & Somersalo, E. Quantitative in silico analysis of neurotransmitter pathways under steady state conditions. Frontiers in endocrinology 4, 137 (2013).

[25] Jolivet, R., Coggan, J. S., Allaman, I. & Magistretti, P. J. Multi-timescale modeling of activity-dependent metabolic coupling in the neuron-glia-vasculature ensemble. PLoS computational biology 11, e1004036 (2015).

[26] Shichkova, P. et al. Breakdown and repair of metabolism in the aging brain. Frontiers in Science 3, 1441297 (2025).

[27] Sertbas, M. & Ulgen, K. O. Exploring human brain metabolism via genomescale metabolic modeling with highlights on multiple sclerosis. ACS Chemical Neuroscience 16, 1346–1360 (2025).

[28] Dienel, G. A. Metabolomic and imaging mass spectrometric assays of labile brain metabolites: Critical importance of brain harvest procedures. Neurochemical Research 45, 2586–2606 (2020).

[29] Rothman, D. L. et al. Glucose sparing by glycogenolysis (gsg) determines the relationship between brain metabolism and neurotransmission. Journal of Cerebral Blood Flow & Metabolism 42, 844–860 (2022).

[30] Sibson, N. R. et al. Stoichiometric coupling of brain glucose metabolism and glutamatergic neuronal activity. Proceedings of the National Academy of Sciences 95, 316–321 (1998).

[31] Rothman, D. L. et al. In vivo 13c and 1h-[13c] mrs studies of neuroenergetics and neurotransmitter cycling, applications to neurological and psychiatric disease and brain cancer. NMR in Biomedicine 32, e4172 (2019).

[32] Rothman, D. L., Behar, K. L. & Dienel, G. A. Mechanistic stoichiometric relationship between the rates of neurotransmission and neuronal glucose oxidation: Reevaluation of and alternatives to the pseudo-malate-aspartate shuttle model. Journal of Neurochemistry 168, 555–591 (2024).

[33] Courchesne-Loyer, A. et al. Inverse relationship between brain glucose and ketone metabolism in adults during short-term moderate dietary ketosis: a dual tracer quantitative positron emission tomography study. Journal of Cerebral Blood Flow & Metabolism 37, 2485–2493 (2017).

[34] Svart, M. et al. Regional cerebral effects of ketone body infusion with 3-hydroxybutyrate in humans: reduced glucose uptake, unchanged oxygen consumption and increased blood flow by positron emission tomography. a randomized, controlled trial. PLoS One 13, e0190556 (2018).

[35] Hertz, L., Dringen, R., Schousboe, A. & Robinson, S. R. Astrocytes: glutamate producers for neurons. Journal of neuroscience research 57, 417–428 (1999).

[36] Rothman, D. L., De Feyter, H. M., de Graaf, R. A., Mason, G. F. & Behar, K. L. 13c mrs studies of neuroenergetics and neurotransmitter cycling in humans. NMR in Biomedicine 24, 943–957 (2011).

[37] Fell, D. & Cornish-Bowden, A. Understanding the control of metabolism Vol. 2 (Portland press London, 1997).

[38] Shepherd, G. M. Neurobiology (Oxford University Press, 1988).

[39] Hertz, L., Peng, L. & Dienel, G. A. Energy metabolism in astrocytes: high rate of oxidative metabolism and spatiotemporal dependence on glycolysis/glycogenolysis. Journal of Cerebral Blood Flow & Metabolism 27, 219–249 (2007).

[40] Dienel, G. A. & Rothman, D. L. Reevaluation of astrocyte-neuron energy metabolism with astrocyte volume fraction correction: impact on cellular glucose oxidation rates, glutamate–glutamine cycle energetics, glycogen levels and utilization rates vs. exercising muscle, and na+/k+ pumping rates. Neurochemical Research 45, 2607–2630 (2020).

[41] Hendry, S. H., Schwark, H., Jones, E. & Yan, J. Numbers and proportions of gaba-immunoreactive neurons in different areas of monkey cerebral cortex. Journal of neuroscience 7, 1503–1519 (1987).

[42] Hyder, F., Fulbright, R. K., Shulman, R. G. & Rothman, D. L. Glutamatergic function in the resting awake human brain is supported by uniformly high oxidative energy. Journal of Cerebral Blood Flow & Metabolism 33, 339–347 (2013).

[43] Blazey, T., Snyder, A. Z., Goyal, M. S., Vlassenko, A. G. & Raichle, M. E. A systematic meta-analysis of oxygen-to-glucose and oxygen-to-carbohydrate ratios in the resting human brain. PLoS One 13, e0204242 (2018).

[44] Eriksen, J., Li, F. & Edwards, R. H. The mechanism and regulation of vesicular glutamate transport: Coordination with the synaptic vesicle cycle. Biochimica et Biophysica Acta (BBA)-Biomembranes 1862, 183259 (2020).

[45] Mason, G. F., Petersen, K. F., De Graaf, R. A., Shulman, G. I. & Rothman, D. L. Measurements of the anaplerotic rate in the human cerebral cortex using 13c magnetic resonance spectroscopy and [1-13c] and [2-13c] glucose. Journal of neurochemistry 100, 73–86 (2007).

[46] Schousboe, A. Transport and metabolism of glutamate and gaba in neurons and glial cells. International review of neurobiology 22, 1–45 (1981).

[47] Mangia, S., Giove, F. & DiNuzzo, M. Metabolic pathways and activity-dependent modulation of glutamate concentration in the human brain. Neurochemical research 37, 2554–2561 (2012).

[48] Soghomonian, J.-J. & Martin, D. L. Two isoforms of glutamate decarboxylase: why? Trends in pharmacological sciences 19, 500–505 (1998).

[49] Waagepetersen, H. S., Sonnewald, U., Gegelashvili, G., Larsson, O. M. & Schousboe, A. Metabolic distinction between vesicular and cytosolic gaba in cultured gabaergic neurons using 13c magnetic resonance spectroscopy. Journal of neuroscience research 63, 347–355 (2001).

[50] Patel, A. B., de Graaf, R. A., Martin, D. L., Battaglioli, G. & Behar, K. L. Evidence that gad65 mediates increased gaba synthesis during intense neuronal activity in vivo. Journal of neurochemistry 97, 385–396 (2006).

[51] Ke, Y., Cohen, B. M., Bang, J. Y., Yang, M. & Renshaw, P. F. Assessment of gaba concentration in human brain using two-dimensional proton magnetic resonance spectroscopy. Psychiatry Research: Neuroimaging 100, 169–178 (2000).

[52] Petroff, O. A., Rothman, D. L., Behar, K. L. & Mattson, R. H. Low brain gaba level is associated with poor seizure control. Annals of Neurology: Official Journal of the American Neurological Association and the Child Neurology Society 40, 908–911 (1996).

[53] Landheer, K., Prinsen, H., Petroff, O. A., Rothman, D. L. & Juchem, C. Elevated homocarnosine and gaba in subject on isoniazid as assessed through 1h mrs at 7t. Analytical biochemistry 599, 113738 (2020).

[54] Fonnum, F. & Walberg, F. The concentration of gaba within inhibitory nerve terminals. Brain research 62, 577–579 (1973).

[55] Alonso-Nanclares, L., Gonzalez-Soriano, J., Rodriguez, J. & DeFelipe, J. Gender differences in human cortical synaptic density. Proceedings of the National Academy of Sciences 105, 14615–14619 (2008).

[56] Rizzoli, S. O. & Betz, W. J. Synaptic vesicle pools. Nature Reviews Neuroscience 6, 57–69 (2005).

[57] Qu, L., Akbergenova, Y., Hu, Y. & Schikorski, T. Synapse-to-synapse variation in mean synaptic vesicle size and its relationship with synaptic morphology and function. Journal of Comparative Neurology 514, 343–352 (2009).

[58] Moulder, K. L. et al. Vesicle pool heterogeneity at hippocampal glutamate and gaba synapses. Journal of Neuroscience 27, 9846–9854 (2007).

[59] Abdallah, C. G. et al. Glutamate metabolism in major depressive disorder. American Journal of Psychiatry 171, 1320–1327 (2014).

[60] Mason, G. F. & Rothman, D. L. Basic principles of metabolic modeling of nmr 13c isotopic turnover to determine rates of brain metabolism in vivo. Metabolic engineering 6, 75–84 (2004).

[61] Buss, K., Drewke, C., Lohmann, S., Piwonska, A. & Leistner, E. Properties and interaction of heterologously expressed glutamate decarboxylase isoenzymes gad65kda and gad67kda from human brain with ginkgotoxin and its 5 ‘-phosphate. Journal of medicinal chemistry 44, 3166–3174 (2001).

[62] Spink, D. C., Porter, T. G., Wu, S. & Martin, D. L. Characterization of three kinetically distinct forms of glutamate decarboxylase from pig brain. Biochemical Journal 231, 695–703 (1985).

[63] Blindermann, J.-M., Maitre, M., Ossola, L. & Mandel, P. Purification and some properties of l-glutamate decarboxylase from human brain. European Journal of Biochemistry 86, 143–152 (1978).

[64] Patel, A., Johnson, A. & Balazs, R. Metabolic compartmentation of glutamate associated with the formation of γ-aminobutyrate. Journal of Neurochemistry 23, 1271–1279 (1974).

[65] Olsen, R. & DeLorey, T. Gaba synthesis, uptake and release. Basic neurochemistry: molecular, cellular and medical aspects 336 (1999).

[66] Mason, G. F. et al. Decrease in gaba synthesis rate in rat cortex following gaba-transaminase inhibition correlates with the decrease in gad67 protein. Brain research 914, 81–91 (2001).

[67] de Graaf, R. A., Patel, A. B., Rothman, D. L. & Behar, K. L. Acute regulation of steady-state gaba levels following gaba-transaminase inhibition in rat cerebral cortex. Neurochemistry international 48, 508–514 (2006).

[68] Schousboe, A., Wu, J. & Roberts, E. Purification and characterization of the 4-aminobutyrate-2-ketoglutarate transaminase from mouse brain. Biochemistry 12, 2868–2873 (1973).

[69] Choi, S. Y. et al. Purification and properties of gaba transaminase from bovine brain. Molecules and Cells 3, 397–401 (1993).

[70] Larsson, O. & Schousboe, A. Kinetic characterization of gaba-transaminase from cultured neurons and astrocytes. Neurochemical research 15, 1073–1077 (1990).

[71] Cooper, A. J. The role of glutamine synthetase and glutamate dehydrogenase in cerebral ammonia homeostasis. Neurochemical research 37, 2439–2455 (2012).

[72] Lebon, V. et al. Astroglial contribution to brain energy metabolism in humans revealed by 13c nuclear magnetic resonance spectroscopy: elucidation of the dominant pathway for neurotransmitter glutamate repletion and measurement of astrocytic oxidative metabolism. Journal of Neuroscience 22, 1523–1531 (2002).

[73] Cooper, A., McDonald, J., Gelbard, A., Gledhill, R. & Duffy, T. The metabolic fate of 13n-labeled ammonia in rat brain. Journal of Biological Chemistry 254, 4982–4992 (1979).

[74] Cooper, A. J. & Jeitner, T. M. Central role of glutamate metabolism in the maintenance of nitrogen homeostasis in normal and hyperammonemic brain. Biomolecules 6, 16 (2016).

[75] Shashidharan, P., Clarke, D. D., Ahmed, N., Moschonas, N. & Plaitakis, A. Nerve tissue-specific human glutamate dehydrogenase that is thermolabile and highly regulated by adp. Journal of neurochemistry 68, 1804–1811 (1997).

[76] Wang, X.-G. & Engel, P. C. Positive cooperativity with hill coefficients of up to 6 in the glutamate concentration dependence of steady-state reaction rates measured with clostridial glutamate dehydrogenase and the mutant a163g at high ph. Biochemistry 34, 11417–11422 (1995).

[77] Patel, A. B. et al. Cerebral pyruvate carboxylase flux is unaltered during bicuculline-seizures. Journal of Neuroscience Research 79, 128–138 (2005).

[78] Gruetter, R., Seaquist, E. R. & Ugurbil, K. A mathematical model of compartmentalized neurotransmitter metabolism in the human brain. American Journal of Physiology-Endocrinology And Metabolism 281, E100–E112 (2001).

[79] Lai, M. et al. In vivo 13c mrs in the mouse brain at 14.1 tesla and metabolic flux quantification under infusion of [1, 6-13c2] glucose. Journal of Cerebral Blood Flow & Metabolism 38, 1701–1714 (2018).

[80] Hassel, B., Westergaard, N., Schousboe, A. & Fonnum, F. Metabolic differences between primary cultures of astrocytes and neurons from cerebellum and cerebral cortex. effects of fluorocitrate. Neurochemical research 20, 413–420 (1995).

[81] Bröer, A. et al. The astroglial asct2 amino acid transporter as a mediator of glutamine efflux. Journal of neurochemistry 73 (1999).

[82] Richards, D. A., Tolias, C. M., Sgouros, S. & Bowery, N. G. Extracellular glutamine to glutamate ratio may predict outcome in the injured brain: a clinical microdialysis study in children. Pharmacological research 48, 101–109 (2003).

[83] Sibson, N. R. et al. In vivo 13c nmr measurements of cerebral glutamine synthesis as evidence for glutamate–glutamine cycling. Proceedings of the National Academy of Sciences 94, 2699–2704 (1997).

[84] Blaauw, R., Nel, D. G. & Schleicher, G. K. Plasma glutamine levels in relation to intensive care unit patient outcome. Nutrients 12, 402 (2020).

[85] Bagga, P. et al. Characterization of cerebral glutamine uptake from blood in the mouse brain: implications for metabolic modeling of 13c nmr data. Journal of Cerebral Blood Flow & Metabolism 34, 1666–1672 (2014).

[86] Boumezbeur, F. et al. The contribution of blood lactate to brain energy metabolism in humans measured by dynamic 13c nuclear magnetic resonance spectroscopy. Journal of Neuroscience 30, 13983–13991 (2010).

[87] Chowdhury, G. M., Jiang, L., Rothman, D. L. & Behar, K. L. The contribution of ketone bodies to basal and activity-dependent neuronal oxidation in vivo. Journal of Cerebral Blood Flow & Metabolism 34, 1233–1242 (2014).

[88] Pan, J. W., Rothman, D. L., Behar, K. L., Stein, D. T. & Hetherington, H. P. Human brain β-hydroxybutyrate and lactate increase in fasting-induced ketosis. Journal of Cerebral Blood Flow & Metabolism 20, 1502–1507 (2000).

[89] Hasselbalch, S. G. et al. Changes in cerebral blood flow and carbohydrate metabolism during acute hyperketonemia. American Journal of Physiology-Endocrinology and Metabolism 270, E746–E751 (1996).

[90] Loman, T. E. et al. Catalyst: Fast and flexible modeling of reaction networks. PLOS Computational Biology 19, e1011530 (2023).

[91] Rackauckas, C. & Nie, Q. Differentialequations. jl–a performant and feature-rich ecosystem for solving differential equations in julia. Journal of open research software 5, 15–15 (2017).

[92] Clarke, K. et al. Kinetics, safety and tolerability of (r)-3-hydroxybutyl (r)-3-hydroxybutyrate in healthy adult subjects. Regulatory Toxicology and Pharmacology 63, 401–408 (2012).

[93] Martin-Arrowsmith, P. W., Lov, J., Dai, J., Morais, J. A. & Churchward-Venne, T. A. Ketone monoester supplementation does not expedite the recovery of indices of muscle damage after eccentric exercise. Frontiers in Nutrition 7, 607299 (2020).

[94] Hannaian, S. J. et al. Acute ingestion of a ketone monoester, whey protein, or their co-ingestion in the overnight postabsorptive state elicit a similar stimulation of myofibrillar protein synthesis rates in young males: a double-blind randomized trial. The American Journal of Clinical Nutrition 119, 716–729 (2024).

[95] Monteyne, A. J. et al. A ketone monoester drink reduces postprandial blood glucose concentrations in adults with type 2 diabetes: a randomised controlled trial. Diabetologia 67, 1107–1113 (2024).

[96] Mujica-Parodi, L. R. et al. Diet modulates brain network stability, a biomarker for brain aging, in young adults. Proceedings of the National Academy of Sciences 117, 6170–6177 (2020).

[97] Fell, D. A. Metabolic control analysis: a survey of its theoretical and experimental development. Biochemical Journal 286, 313 (1992).

[98] Flanagan, B., McDaid, L., Wade, J., Wong-Lin, K. & Harkin, J. A computational study of astrocytic glutamate influence on post-synaptic neuronal excitability. PLoS computational biology 14, e1006040 (2018).

[99] Cunnane, S. C. et al. Can ketones compensate for deteriorating brain glucose uptake during aging? implications for the risk and treatment of alzheimer’s disease. Annals of the New York Academy of Sciences 1367, 12–20 (2016).

[100] Lund, T. M., Risa, Ø., Sonnewald, U., Schousboe, A. & Waagepetersen, H. S. Availability of neurotransmitter glutamate is diminished when β-hydroxybutyrate replaces glucose in cultured neurons. Journal of neurochemistry 110, 80–91 (2009).

[101] Owen, O. et al. Brain metabolism during fasting. The Journal of clinical investigation 46, 1589–1595 (1967).

[102] Petroff, O. A. & Rothman, D. L. Measuring human brain gaba in vivo: effects of gaba-transaminase inhibition with vigabatrin. Molecular neurobiology 16, 97–121 (1998).

[103] Petroff, O. A., Behar, K. L., Mattson, R. H. & Rothman, D. L. Human brain γ-aminobutyric acid levels and seizure control following initiation of vigabatrin therapy. Journal of neurochemistry 67, 2399–2404 (1996).

[104] Walls, A. B. et al. Knockout of gad65 has major impact on synaptic gaba synthesized from astrocyte-derived glutamine. Journal of Cerebral Blood Flow & Metabolism 31, 494–503 (2011).

[105] Buddhala, C., Wei, J. & Wu, J. Y. Activity dependent cleavage and protein phosphorylation regulate human glutamate decarboxylase gad65 activity (2008).

[106] Bhagwagar, Z. et al. Increased brain gaba concentrations following acute administration of a selective serotonin reuptake inhibitor. American Journal of Psychiatry 161, 368–370 (2004).

[107] Ebrahimi Behnam, B., Raeisi, S., Bakhtiari, N., Abbasi, M. M. & Shanehbandi, D. Neurochemical and genetic effects of the ketogenic diet: alterations in brain gaba, glutamate, and related gene. Molecular Biology Reports 52, 968 (2025).

[108] Shulman, R. & Rothman, D. Enzymatic phosphorylation of muscle glycogen synthase: a mechanism for maintenance of metabolic homeostasis. Proceedings of the National Academy of Sciences 93, 7491–7495 (1996).

[109] Rothman, D. L., Stearns, S. C. & Shulman, R. G. Gene expression regulates metabolite homeostasis during the crabtree effect: implications for the adaptation and evolution of metabolism. Proceedings of the National Academy of Sciences 118, e2014013118 (2021).

[110] Rothman, D. L., Moore, P. B. & Shulman, R. G. The impact of metabolism on the adaptation of organisms to environmental change. Frontiers in Cell and Developmental Biology 11, 1197226 (2023).

[111] McKenna, M. C. Substrate competition studies demonstrate oxidative metabolism of glucose, glutamate, glutamine, lactate and 3-hydroxybutyrate in cortical astrocytes from rat brain. Neurochemical research 37, 2613–2626 (2012).

[112] Terpstra, M. et al. Changes in human brain glutamate concentration during hypoglycemia: insights into cerebral adaptations in hypoglycemia-associated autonomic failure in type 1 diabetes. Journal of Cerebral Blood Flow & Metabolism 34, 876–882 (2014).

[113] Owen, O. E., Kalhan, S. C. & Hanson, R. W. The key role of anaplerosis and cataplerosis for citric acid cycle function. Journal of Biological Chemistry 277, 30409–30412 (2002).

[114] Ma, W., Berg, J. & Yellen, G. Ketogenic diet metabolites reduce firing in central neurons by opening katp channels. Journal of Neuroscience 27, 3618–3625 (2007).

[115] Bough, K. J. & Rho, J. M. Anticonvulsant mechanisms of the ketogenic diet. Epilepsia 48, 43–58 (2007).

[116] Dienel, G. A. & Cruz, N. F. Contributions of glycogen to astrocytic energetics during brain activation. Metabolic brain disease 30, 281–298 (2015).

[117] Hijazi, S., Smit, A. B. & van Kesteren, R. E. Fast-spiking parvalbumin-positive interneurons in brain physiology and alzheimer’s disease. Molecular Psychiatry 28, 4954–4967 (2023).

[118] Small, S. et al. The pathogenicity of the glutamate metabolic cycle in schizophrenia determined by hippocampal spectroscopic profiling. Research Square (2025).

[119] Uno, Y. & Coyle, J. T. Glutamate hypothesis in schizophrenia. Psychiatry and clinical neurosciences 73, 204–215 (2019).

[120] Gigante, A. D. et al. Brain glutamate levels measured by magnetic resonance spectroscopy in patients with bipolar disorder: a meta-analysis. Bipolar disorders 14, 478–487 (2012).

[121] Schwartz, T. L., Nihalani, N., Jindal, S., Virk, S. & Jones, N. Psychiatric medication-induced obesity: a review. Obesity reviews 5, 115–121 (2004).

[122] Hilt, R. J. et al. Side effects from use of one or more psychiatric medications in a population-based sample of children and adolescents. Journal of child and adolescent psychopharmacology 24, 83–89 (2014).

[123] Lehallier, B. et al. Undulating changes in human plasma proteome profiles across the lifespan. Nature medicine 25, 1843–1850 (2019).

[124] Shen, X. et al. Nonlinear dynamics of multi-omics profiles during human aging. Nature aging 1–16 (2024).

[125] Wierenga, C. E. & Bondi, M. W. Use of functional magnetic resonance imaging in the early identification of alzheimer’s disease. Neuropsychology review 17, 127–143 (2007).

[126] Sperling, R. The potential of functional mri as a biomarker in early alzheimer’s disease. Neurobiology of aging 32, S37–S43 (2011).

[127] Wang, H. E. et al. Virtual brain twins: from basic neuroscience to clinical use. National Science Review 11, nwae079 (2024).

[128] Hofmann, D. et al. Increasing spectral dcm flexibility and speed by leveraging julia’s modelingtoolkit and automated differentiation. bioRxiv 2023–10 (2024).

[129] Pathak, A. et al. Biomimetic model of corticostriatal micro-assemblies discovers a neural code. Nature Communications (2025).

