## Supplemental Information for "Computational modeling of neurotransmitter cycling predicts human brain glutamate and GABA dynamics in response to administration of exogenous ketones"

**Table 1** Datasets used for estimates of compartmental concentrations.

| Metabolite pool | Value | Reference |
|---|---|---|
| $[Glu]_e$ | 9.8 ± 2.4 $\mu$mol/g | [1] |
| $[Glu]_e$ | 5-15 $\mu$mol/g | [2] |
| $[Glu]_e$ | 10-15 mM | [3] |
| $[Glu]_a$ | 0.8 $\mu$mol/g | [1] |
| $[Glu]_i$ | 2% of total glutamate | [4] |
| $[GABA]_{total}$ | 1.09 ± 0.4 $\mu$mol/cm$^3$ | [5] |
| $[GABA]_{total}$ | 1.18 ± 0.06 mmol/kg | [6] |
| $[GLN]_{total}$ | 4.2 ± 1.2 $\mu$mol/g | [1] |
| $[GLN]_a$ | ∼90% of total glutamine | [7] |
| $[GLN]_{ecf}$ | 249-572 $\mu$mol/L | [8] |
| $[GLN]_{ecf}$ | 0.13-0.5 mM | [9] |
| $[GLN]_p$ | 420-700 $\mu$mol/L | [10] |

**Table 2** Datapoints used to estimate ketone metabolism related parameters.

| Reference | $\mathbf{[Ket]_p}$ | $\mathbf{[Ket]_b}$ | $\mathbf{CMR_{ket}}$ |
|---|---|---|---|
| Pan et al. 2002 [11] | 2.25 mM | 0.18 mM | 0.032 mM/min |
| Svart et al. 2018 [12] | 5.5 mM | 0.44 mM* | 0.070 mM/min† |

* Calculated from $[Ket]_p$ and assuming linear scaling with datapoint from Pan et al. 2002

† Calculated from $\Delta\%$ $CMR_{glc}$ = 14%

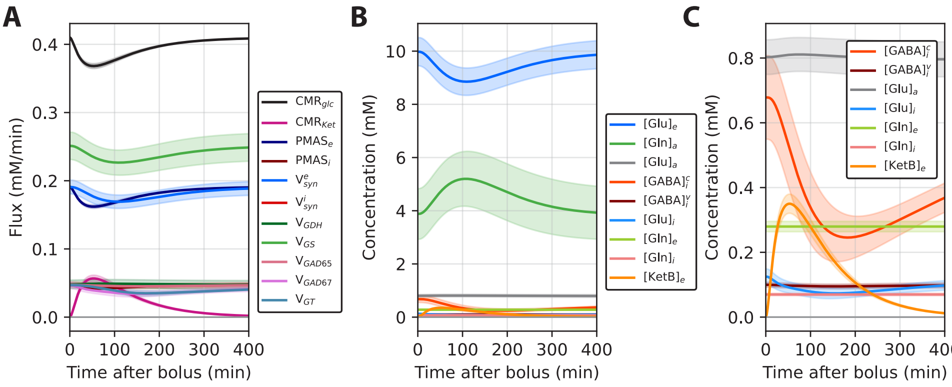


**Fig. 1 Simulated time courses of neurotransmitter cycling fluxes and concentrations, scaled to total tissue volume. A**: Time courses of fluxes after a D-$\beta$HB bolus. Error bands represent the standard deviation obtained by varying parameters by ±10% around their central values. **B**: Time courses of absolute concentrations. **C**: Time courses of absolute concentrations shown on a smaller scale to resolve changes in the smaller metabolite pools.

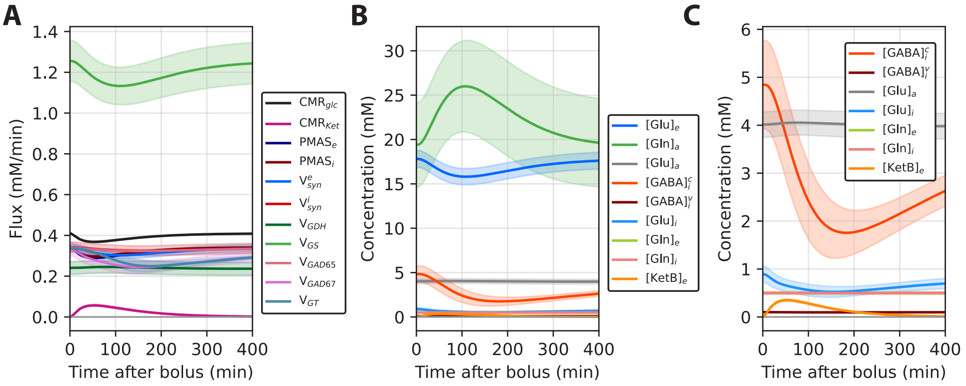


**Fig. 2 Simulated time courses of neurotransmitter cycling fluxes and concentrations, scaled to compartmental volumes. A**: Time courses of fluxes after a D-$\beta$HB bolus. Error bands represent the standard deviation obtained by varying parameters by $\pm 10\%$ around their central values. **B**: Time courses of absolute concentrations. **C**: Time courses of absolute concentrations shown on a smaller scale to resolve changes in the smaller metabolite pools.

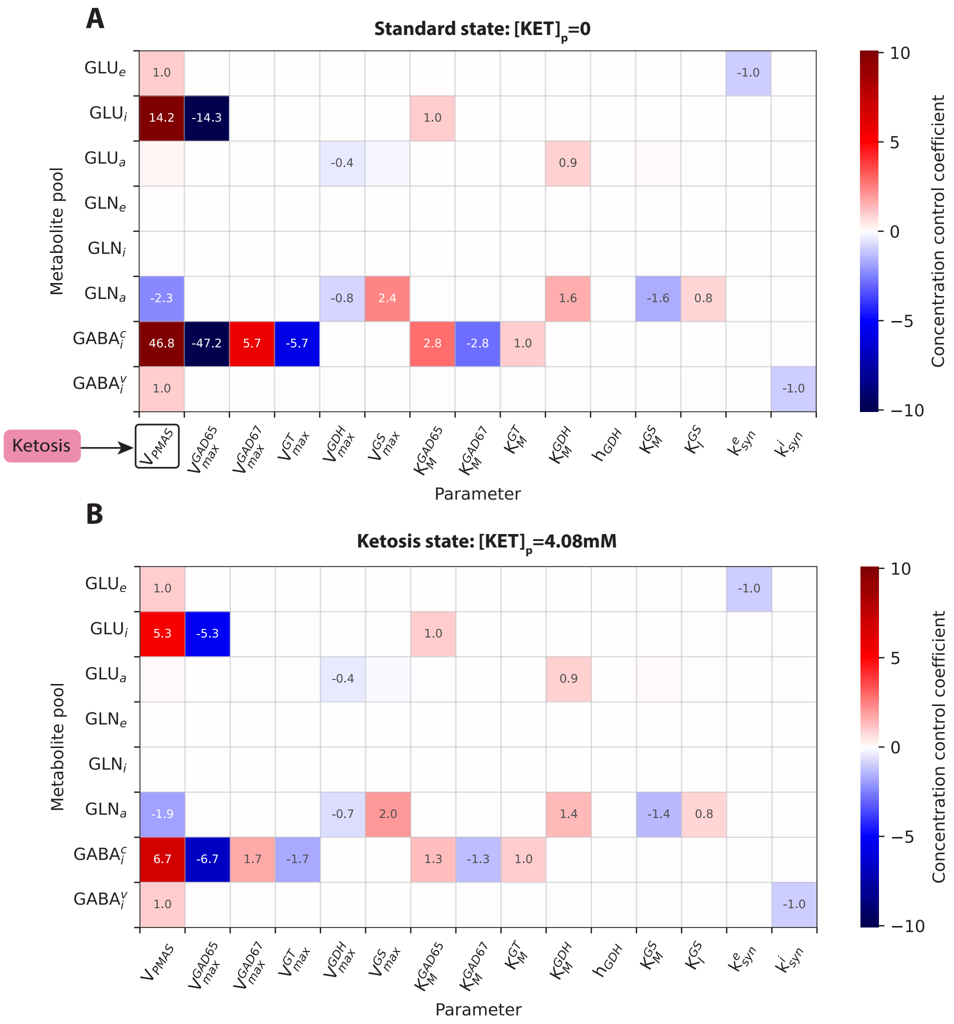


**Fig. 3 Control coefficients from metabolic control analysis of glutamate and GABA concentrations in neurotransmitter cycling across different conditions. A**: Standard state (no ketones). **B**: Ketosis state ($[Ket]_p$ fixed at 4.08 mM to maintain steady state). Notably, control patterns were preserved across conditions, with differences observed only in magnitude.

### References


[1] Mason, G. F., Petersen, K. F., De Graaf, R. A., Shulman, G. I. & Rothman, D. L. Measurements of the anaplerotic rate in the human cerebral cortex using 13c magnetic resonance spectroscopy and [1-13c] and [2-13c] glucose. *Journal of neurochemistry* **100**, 73–86 (2007).

[2] Schousboe, A. Transport and metabolism of glutamate and gaba in neurons and glial cells. *International review of neurobiology* **22**, 1–45 (1981).

[3] Mangia, S., Giove, F. & DiNuzzo, M. Metabolic pathways and activity-dependent modulation of glutamate concentration in the human brain. *Neurochemical research* **37**, 2554–2561 (2012).

[4] Patel, A., Johnson, A. & Balazs, R. Metabolic compartmentation of glutamate associated with the formation of $\gamma$-aminobutyrate. *Journal of Neurochemistry* **23**, 1271–1279 (1974).

[5] Ke, Y., Cohen, B. M., Bang, J. Y., Yang, M. & Renshaw, P. F. Assessment of gaba concentration in human brain using two-dimensional proton magnetic resonance spectroscopy. *Psychiatry Research: Neuroimaging* **100**, 169–178 (2000).

[6] Petroff, O. A., Rothman, D. L., Behar, K. L. & Mattson, R. H. Low brain gaba level is associated with poor seizure control. *Annals of Neurology: Official Journal of the American Neurological Association and the Child Neurology Society* **40**, 908–911 (1996).

[7] Hassel, B., Westergaard, N., Schousboe, A. & Fonnum, F. Metabolic differences between primary cultures of astrocytes and neurons from cerebellum and cerebral cortex. effects of fluorocitrate. *Neurochemical research* **20**, 413–420 (1995).

[8] Richards, D. A., Tolias, C. M., Sgouros, S. & Bowery, N. G. Extracellular glutamine to glutamate ratio may predict outcome in the injured brain: a clinical microdialysis study in children. *Pharmacological research* **48**, 101–109 (2003).

[9] Bröer, S. & Brookes, N. Transfer of glutamine between astrocytes and neurons. *Journal of neurochemistry* **77**, 705–719 (2001).

[10] Blaauw, R., Nel, D. G. & Schleicher, G. K. Plasma glutamine levels in relation to intensive care unit patient outcome. *Nutrients* **12**, 402 (2020).

[11] Pan, J. W. *et al.* [2, 4-13c2]-$\beta$-hydroxybutyrate metabolism in human brain. *Journal of Cerebral Blood Flow & Metabolism* **22**, 890–898 (2002).

[12] Svart, M. *et al.* Regional cerebral effects of ketone body infusion with 3-hydroxybutyrate in humans: reduced glucose uptake, unchanged oxygen consumption and increased blood flow by positron emission tomography. a randomized,

controlled trial. *PLoS One* **13**, e0190556 (2018).
